# Hunting the plant surrender signal activating apoplexy in grapevines after *Neofusicoccum parvum* infection

**DOI:** 10.1101/2021.10.14.464072

**Authors:** Islam M. Khattab, Jochen Fischer, Andrzej Kaźmierczak, Eckhard Thines, Peter Nick

## Abstract

- Apoplectic breakdown from Grapevines Trunk Diseases (GTDs) has become a serious challenge to viticulture in consequence to drought stress. We hypothesise that fungal aggressiveness is controlled by a chemical communication between host and colonising fungus.
- We introduce the new concept of a “plant surrender signal” accumulating in host plants under stress and triggering aggressive behaviour of the strain *Neofusicoccum parvum* (Bt-67) causing Botryosphaeriaceae-related dieback in grapevines.
- Using a cell-based experimental system (*Vitis* cells) and bioactivity-guided fractionation, we identify *trans*-ferulic acid, a monolignol precursor, as “surrender signal”. We show that this signal specifically activates secretion of the fungal phytotoxin Fusicoccin A. We show further that this phytotoxin, mediated by 14-3-3 proteins, activates programmed cell death in *Vitis* cells.
- We arrive at a model pinpointing the chemical communication driving apoplexy in *Botryosphaeriaceae-Vitis* interaction and define the channelling of phenylpropanoid pathway from the lignin precursor, *trans*-ferulic acid to the phytoalexin trans-resveratrol as target for future therapy.

## 1. Introduction

Concomitantly with climate change, *Botryosphaeriaceae*-related Dieback turned into a progressively devastating threat for viticulture. Already a decade ago, the economic damage by Grapevine Trunk Diseases (GTDs) was estimated to exceed 1500 million US$ per year (Hofstetter *et al*., 2012); for France alone yield was reduced by 25% or ~ 5000 million US$ (https://www.maladie-du-bois-vigne.fr) in 2016. The interaction of *Botryosphaeriaceae* with their grapevine hosts is very complex. They live as endophytes for many years during a latent phase but can suddenly switch to an apoplectic phase killing the grapevine host within a few days (Slippers & Wingfield, 2007). This lifestyle typical of GTDs differs from other pathogens because it does not meet Koch’s postulates, since the presence of a pathogen is sufficient to cause host symptoms (Loeffler F. 1884). During the endophytic phase of wood-decaying fungi this link seems to be absent. It is a change of fungal behaviour, not the presence of the microbe that causes apoplexy (**Fig. 1**).

**Fig. 1.**
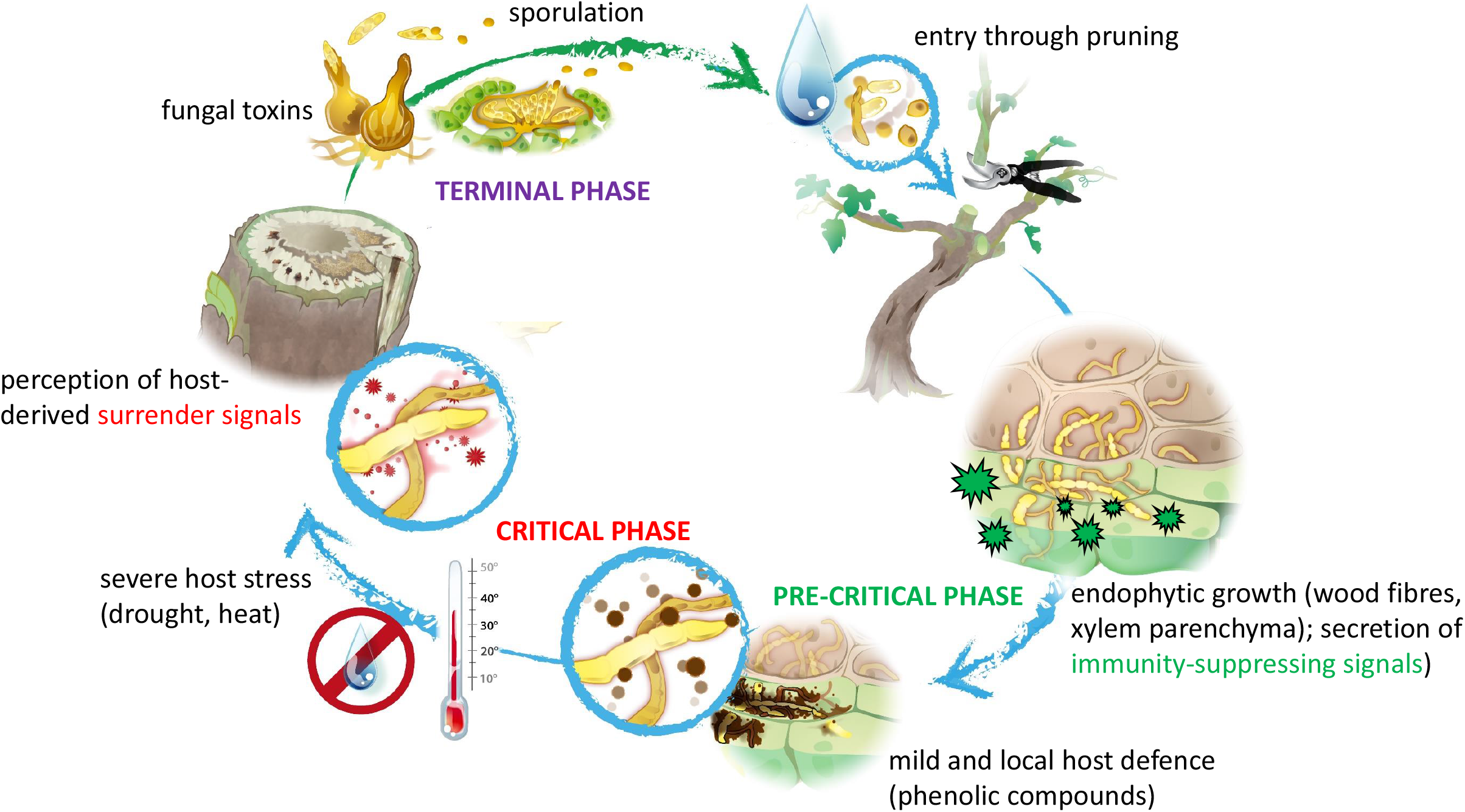
Working model on host-pathogen interaction during *Vitis-Botryosphaeriaceae* Dieback used to structure the current study. After entry through pruning wounds, the fungus first colonises wood fibres and xylem parenchyma and is contained by a local and mild defence response of the host (accumulation of phenolic compounds). Host defence is partially quelled by immunity-suppressing signals secreted by the endophyte. This pre-critical phase can last for years and under favourable conditions will not impair productivity nor vigour of the host. However, if the host is shifted under severe stress such as heat or drought, the critical phase can be initiated. Here, the stress-dependent, chronic loss of physiological homeostasis of the host results in molecular changes that can be perceived by the fungus as surrender signals and activate a switch from asymptomatic endophytic growth to active killing of the host. This critical phase is, therefore, followed by a terminal phase, where the fungus is producing toxins and initiates sporulation and formation of fruiting bodies resulting in the death of the host and the spread of spores that can colonise neighbouring vines.

GTDs have always been present in Europe. The first report, in the book *Kitab Al-Felaha* by the Andalusian Scientist *Ibn Al-Awam*, dates back to the twelfth century (Mugnai *et al*., 1999). However, outbreaks have turned more common with climate change (Slippers & Wingfield, 2007; Galarneau *et al*., 2019). Thus, GTDs clearly differ from classical diseases, such as Downy Mildew or Powdery Mildew, as diseases with the largest impact on viticulture (Jürges *et al*., 2009). Although GTD has been manifest for a long time, we know very little about the molecular mechanisms that provoke a harmless endophyte to turn into a dangerous, toxin-producing killer. Although the impact of plant-pathogen crosstalk in the context of GTDs is clear, we are still far from understanding this crosstalk.

Among the numerous species of the *Botryosphaeriaceae* associated with dieback in grapevines (Carlucci *et al*., 2015), the aggressive species *Neofusicoccum parvum* has served as experimental model (Úrbez-Torres & Gubler, 2009; Stempien *et al*., 2017).

Colonisation begins from wounded-wood tissues, usually in the context of pruning (Djoukeng *et al*., 2009; Úrbez-Torres, 2011), and spreads through xylem vessels or parenchymatic rays (Gómez *et al*., 2016; Massonnet *et al*., 2017; Khattab *et al*., 2021).

The apoplectic breakdown of the host plant seems to result from fungal phytotoxins reaching the leaves with the transpiration stream, along with hydraulic failure by tyloses and gelous depositions in the infected vessels (Bortolami *et al*., 2019). While wider xylem vessels were proposed to promote susceptibility against *Phaeomoniella* (Pouzoulet *et al*., 2017), a comparative study for *N. parvum* did not show any correlation of fungal spread with vessel diameter, but rather with differential channelling of the stilbene pathway (Khattab *et al*., 2021). A further hallmark for susceptibility are key genes of lignin biosynthesis, i.e.,*caffeic acid O-methyltransferase* channelling metabolism towards lignin (Khattab *et al*., 2021).

The molecular mechanisms behind apoplectic breakdown are still elusive. It is clear, however, that the outbreak of GTDs is promoted by drought stress, as a consequence of global warming, which holds true for both, the Esca syndrome (Lima et al, 2017), and *Botryosphaeriaceae*-related Dieback (Galarneau et al., 2019). While the presence of the fungus is necessary for disease outbreak, it is not a sufficient condition for apoplexy. Since apoplexy is a conditional phenomenon, it might result from altered chemical communication between plant and fungus (**Fig. 1**). During the endophytic phase, the fungus obviously does not produce the compounds activating apoplexy. The fungus releases those compounds conditionally, i.e., when the host shifts to drought stress. The selective advantage of this behavioural switch might be the need to form a fruiting body to escape from a dwindling host. The ability to change depending on the well-being of the host means that the fungus must be able to sense the state of the host plant. Conceptually, the conditional release of phytotoxic compounds derives from a “surrender signal” originating from a host under drought stress.

In the present study, we test the hypothesis that the *Botryosphaeriaceae*-related Dieback in grapevines results from a chemical communication between host and pathogen. We used *N. parvum* Bt-67, as one of the most aggressive and virulent fungal strains causing this disease (Guan et al., 2016; Stempien et al., 2017), and a cell-based experimental system to study plant-fungal interaction *in vitro*. This system allowed us to screen candidates for the “surrender signal”, i.e., for plant compounds that accumulate under drought stress and can trigger the release of fungal compounds with phytotoxic activity. We first identified the lignin precursor ferulic acid as specific and efficient activator for the release of fungal phytotoxins. Using bioactivity-guided fractionation, we were then able to identify a Fusicoccin A (FCA) derivative as *bona-fide* candidate for the phytotoxic effect. In the third part of the study, we analysed the mode of action of FCA and found that this compound induced programmed cell-death (PCD) in grapevine cells. Thus, the fungal response to the plant-derived “surrender signal” is a signal itself that can evoke a plant response that under a different functional context is meaningful for the plant. In the context of host-endophyte interaction under external stress, this response is deleterious, however. In other words – the fungus hijacks plant responses for the sake of its own survival.

## 2. Methodology

### Plant and fungal materials

The strain *Neofusicoccum parvum Bt-67*, isolated by Instituto Superior de Agronomia, Universidade de Lisbona, Portugal (Stempien *et al*., 2017), was kindly provided by the Laboratoire Vigne Biotechnologies et Environnement, Université de Haute-Alsace, France. We conducted the experiments with the grapevine suspension line Vrup-TuB6-GFP, deriving from *Vitis rupestris*, and expressing a GFP fused with ß-tubulin 6 (Guan *et al*., 2015). The response of actin filaments was followed using a cell line in *Vitis vinifera* cv. Chardonnay expressing a Fluorescent fimbrin actin-binding domain 2 (*At*FABD2)-GFP (Akaberi *et al*., 2018). Additionally, two tobacco (*Nicotiana tabacum* L. cv. ‘Bright-Yellow 2’) lines overexpressing *V. rupestris* Metacaspases 2 and 5 were used along with their non-transformed wild type (Gong *et al*., 2019). All cells were subcultured in weekly intervals into Murashige and Skoog medium as described in Akaberi *et al*. (2018) using the appropriate antibiotics. All experiments were conducted with cells that were not cycling, either subsequently to (day 4 in grapevine cells) or prior to the division phase (day 1 for tobacco BY-2 cells).

### Screening the effect of monolignol precursors on *N. parvum* aggressiveness

To probe whether the monolignol precursors cinnamic acid, *p-*coumaric acid, caffeic acid, and *trans*-ferulic acid can induce *N. parvum* to secrete phytotoxic compounds, fungal mycelia were fermented with 0.5, 1, or 1.5 mM for each of these precursors. Fungal mycelia, grown for 2 weeks on Potato Dextrose Agar, PDA (Sigma-Aldrich, Deisenhofen), were cultivated with the respective compound in 250 ml of 20 g^•^L^-1^ malt extract medium (Carl Roth GmbH, Karlsruhe, Germany), pH 5.3 for two additional weeks. After sterile filtration through a 0.22-µ m PVDF membrane (Carl Roth GmbH, Karlsruhe, Germany), the culture filtrate was added to the Vrup-TuB6 cells (35 µl. filtrate/ml cell suspension) to score mortality (see below).

To explore whether phytotoxic compounds might be retained inside the hyphae without being secreted, we extracted fungal metabolites, either from medium or mycelia, after 24 h fermentation with the respective precursor after the fungus had consumed the sugar entirely, which was checked with a Diabur 5000 test strip (Roche, Basel; Switzerland).

### Extraction of fungal metabolites and analysis by HPLC and HPLC-MS

The fungus was cultured in 20 L Yeast Malt Glucose medium (yielding from the culture filtrate about 7.2 g of a crude extract after *trans*-ferulic acid supplement) as explained in **S. Method 1**. Fractions of increasing hydrophobicity (0-100% MeCN) were then tested for their bioactivity in the *Vitis* cell culture system. To identify the fungal phytotoxin released in response to *trans*-ferulic acid, the most toxic phase was re-analysed by a HPLC-MS (Series 1200, Agilent, Waldbronn, Germany) equipped with an UV-DAD and a coupled LC/MSD Trap APCI-mass spectrometer with positive and negative polarisation as described by Buckel et al. (2017). For the mass spectrum of the derived molecules as well as their HPLC-MS analysis see **Supplementary. 2**. The HPLC-MS analysis of the fraction with the strongest phytotoxicity upon *Vitis* cells identified a derivative of Fusicoccin A.

### Mapping the responses to by Fusicoccin A

After identifying Fusicoccin A (FCA) as fungal phytotoxin, we dissected the cellular responses to 6 and 12 µM synthetic FCA (Santa Cruz Biotechnology, Heidelberg, Germany). in Vrup-TuB6-GFP. To get insight into the cellular events involved in this response, a pharmacological strategy was employed as follows:

### Inhibiting Respiratory burst Oxygen Homolog (RBOH)

Prior to FCA treatment, *Vitis* cells were pre-treated for 60 min with 1 µM of Diphenyleneiodonium chloride (DPI; Sigma-Aldrich, Germany) inhibiting NADPH oxidases in the plasma-membrane, diluted in 100 µM DMSO (Chang *et al*., 2011).

### Blocking the FCA receptor, a 14-3-3 protein

Fusicoccin receptors were blocked using pretreatment for 60 min with 5 µM (in 100 µM DMSO) of BV02 (Sigma-Alrich, Germany) in 100 µM DMSO which inhibits 14-3-3 proteins docking sites (Stevers *et al*., 2018).

### Measurement of extracellular pH

We followed changes in the extracellular pH in Vrup-TuB6-GFP cells by a pH meter (Schott handylab, pH12) recording by a digital memory recorder displaying the pH at 1-second intervals as detailed in Qiao *et al*. (2010). To probe the role of 14-3-3 proteins as FCA receptors on ATPases activity, cells were pre-incubated with 5 µM of the 14-3-3 blocker for 45 min, and then treated by FCA (6 µM).

### Superoxide [O_2_^-^] detection

We estimated the intracellular superoxide accumulation in *Vitis* cells by the 0.1% (w/v) Nitroblue Tetrazolium (NBT) assay as described in Steffens and Sauter (2009) and Pietrowska *et al*. (2014) with modifications. After filtering suspension cells from the culture medium, we incubated the *Vitis* cells in NBT for 1 h under aseptic conditions before washing in phosphate buffer prior to observation by bright-field microscopy (Axioskop, Zeiss). Cells accumulating superoxide appeared in blue colour. The frequency of stained cells served as readout for superoxide accumulation, scoring 700 individual cells for each of three biological replicates of the respective time-point.

### Modulating salicylic acid biosynthesis and mitogen-activated protein kinase (MAPK) signalling

To assess the role of salicylic acid in FCA signalling, we pre-treated the *Vitis* cells with 50 µM of salicylic acid (Sigma Aldrich, Deisenhofen, Germany) dissolved in 0.2% (v/v) ethanol for 1 h prior to FCA treatment. Furthermore, we used 1-aminobenzotriazole (ABT; Sigma Aldrich, Deisenhofen, Germany) as an inhibitor for salicylic acid biosynthesis (Leon *et al*., 1995). Here, we incubated the *Vitis* cells with 25 µM of ABT for 1 h prior to FCA treatment. To check whether MAPKs are involved in the signalling deployed by FCA, *Vitis* cells were treated first with 50 µM of the MAPK inhibitor PD98059 (Sigma Aldrich, Germany) as described by Duan *et al*. (2016), prior to FCA treatment.

### Live-cell imaging of the cytoskeleton

We followed the response of the cytoskeleton to FCA using the microtubule marker line Vrup-TuB6 expressing *At*TUB6-GFP (Guan *et al*., 2015), and the actin marker line cv. ‘Chardonnay’ expressing Vv-*At*FABD2-GFP (Akaberi *et al*., 2018). The cytoskeletal response was captured by spinning disc confocal microscopy with an AxioObserver Z1 (Zeiss, Jena, Germany) microscope, equipped with a spinning-disc device (YOKOGAWA CSU-X1 5000), and 488 nm emission light from an Ar-Kr laser (Wang & Nick 2017). We collected confocal z-stacks, processed them by the ZEN software (Zeiss, Oberkochen), and exported them as TIFF format.

To quantify the width of actin filaments as readout for actin bundling, we transformed the images into binary format, adjusting the threshold with the B/W option of the ImageJ freeware (https://imagej.nih.gov/ij). Using the Analyze particles tool, actin filaments were selected automatically, filtering against random signals using a detection threshold of 10 square pixels, and the fit ellipse command to fit the detected particles and to quantify the short ellipse axis as readout for filament width. The integrity of cortical microtubules was estimated according to Schwarzerová, *et al*. (2002). We collected four intensity profiles along the long axis of the cell in equal spacing over the cross axis and using a line width of 25 pixels according to modifications described in Guan *et al*. (2020). For both, microtubules, and actin filaments, twenty-five individual cells were scored for each treatment representing 4 independent biological replicates.

### Manipulation of the cytoskeleton

To assess the role of microtubules on the response to FCA, Vrup-TuB6-GFP were either pre-treated with 10 µM of Taxol (Sigma-Aldrich, Germany), which stabilises microtubules, or with 10 µM of Oryzalin, eliminating microtubules (Sigma-Aldrich, Germany) for 30 min, before adding FCA. To assess the role of actin filaments, cv. Chardonnay Expressing *At*FABD2-GFP cells were pre-treated with 2 µM of the actin-eliminating compound Latrunculin B (Sigma-Aldrich, Deisenhofen, Germany) for 1 h prior to FCA treatment.

### RNA extraction and qPCR

*V. rupestris* (*At*TUB6-GFP) cells were collected and immediately frozen in liquid nitrogen. We extracted total RNA with the RNA Purification Kit (Roboklon, Berlin, Germany), generated cDNA from 1 µg of total RNA, pre-incubating with oligo (dT) primers, followed by reverse transcription with M-MuLV enzyme (New England Biolabs; Frankfurt, Germany) in presence of RNase inhibitor.

We monitored transcripts of genes of interests by qPCR using CFX-PCR System (Bio-Rad, München; Germany) as described in Svyatyna *et al*. (2014). The housekeeping gene, *EF-1α*, served to calculate the steady-state transcripts of target genes using the 2^-ΔΔCt^ method (Livak and Schmittgen, 2001). For details of primer sequences, and accession numbers of target genes see **Table S1**.

### Cell-death assays

To obtain insight into the nature of cell-death, we double-stained the cells for two min with Acridine Orange, AO (100 µg^•^mL^-1^) and Ethidium Bromide, EB (100 µg^•^mL^-1^) simultaneously (Byczkowska *et al*., 2013). The stained cells were analysed by fluorescence microscopy (Diaplan, Leitz) upon excitation in the blue (filter set I3, excitation 450–490 nm, beam splitter 510 nm) recording emission above 515 nm and digital image acquisition (Leica DFC 500 and Leica Application Suite, v4). Living cells exclude EB and appear green. Cells in the 1^st^ stage show penetration of EB only into the cytoplasm, giving greenish yellow to yellow nuclei. Cells in the 2^nd^ stage show bright orange nuclei, because EB crosses the nuclear envelope. Dead cells lose the AO after the breakdown of plasmamembrane, while EB remains sequestered at the DNA, displaying red nuclei. Frequency distributions represent 300-400 individual cells collected from three independent experimental series.

In addition, we used the Evans Blue Dye Assay (Gaff and Okong’o-Ogola, 1971) which labels dead cells blue to a loss of membrane integrity. We stained the *Vitis* cells with 2.5 % Evans blue (Sigma Aldrich, Germany) for 3 min, and then washed twice with distilled water to remove unbound dye. Mortality scores represent populations of at least 1000 individual cells for each biological replicate of the respective treatment.

### The effect of drought on fungal colonisation and lignin accumulation

Disease development was evaluated in infected grapevines under two water-regimes in the variety *V. vinifera* cv. ‘Augster Weiss’. After pre-cultivation of woodcuttings for 3 months in pots in the greenhouse, we infected homogenously individuals with *N. parvum* Bt-67 mycelia according to Khattab *et al*. (2021), and directly subdivided into two experimental sets. One set was well irrigated in 5 days/week and served as control, while the other set developed under constrained irrigation (watering only once weekly). Each set consisted of 3 infected plants and the respective mock control. We evaluated the result 30 days later for wood necrosis and lignin content according to Khattab *et al*. (2021) in the infection site.

## 3. Results

### Drought stress promotes infection development and lignin accumulation

Since drought, as accentuated by climate change, seems to promote disease outbreak, we tested whether restricted irrigation would accentuate the grapevine response to infection with *N. parvum*. The imposed drought stress was sufficient to induce significant symptoms as reduced growth and partial leaf discoloration **(Fig. 2a)**. The necrosis due to *N. parvum* infection (**Fig. 2b**, *NP*) was almost twice in the stressed plants. To check, whether this increase in symptomatic wood was related to fungal development, we used a *Botryosphaeriaceae* specific internal transcribed spacer (*its*) for quantification (**Method. S1: Fig. 1 S1**) and found more than twice fungal DNA under drought stress in the inoculation site itself, while the two sites 3 cm below or above did not differ.

**Fig. 2.**
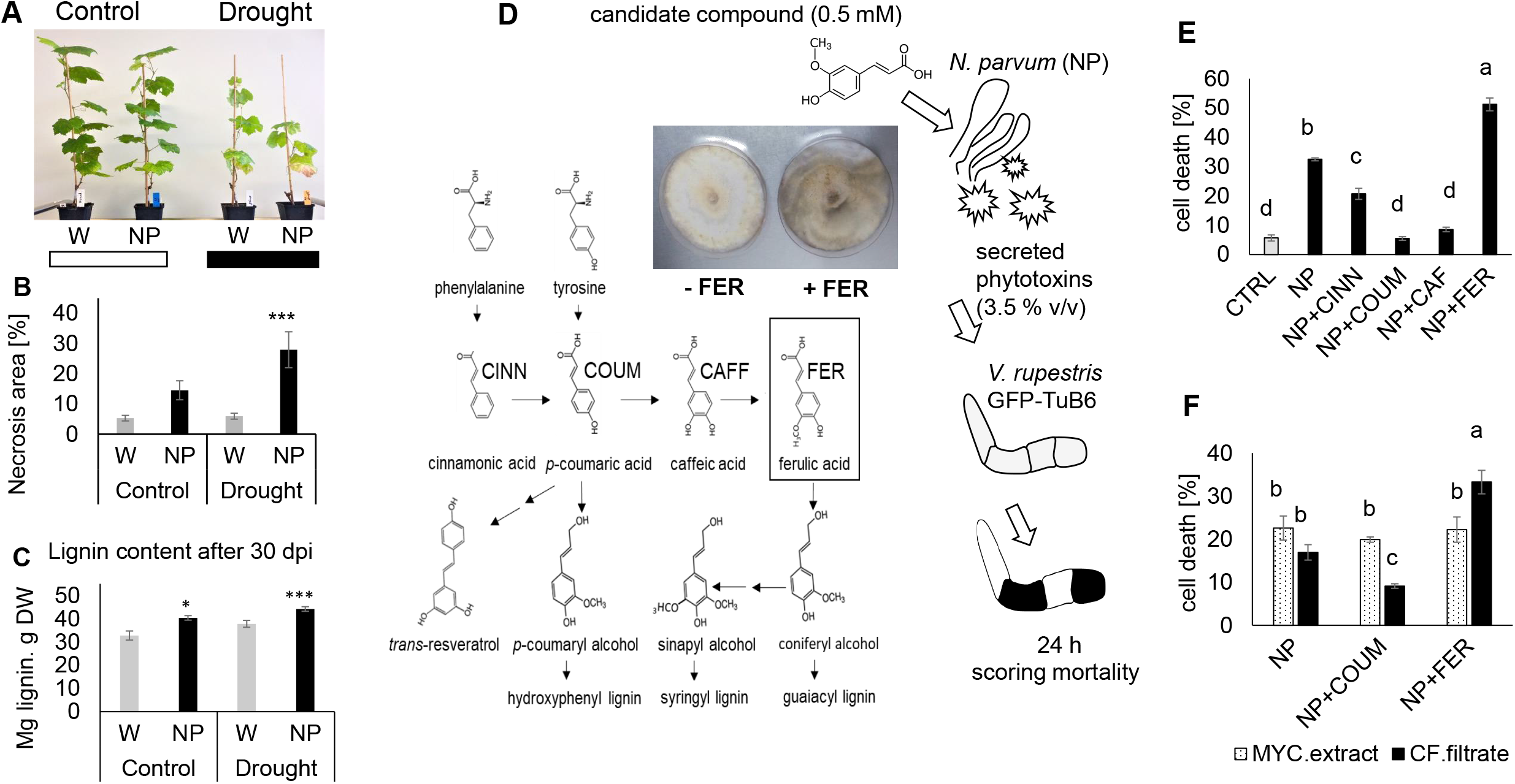
Emerging the fungal strain, *Neofusicoccum parvum* Bt-67, to necrotrophic lifestyle. **A)** Potted grapevines (Augster weiß cultivar) under two experimental sets, either control conditions (optimal water regime) or drought stress (20% of optimal water regime). **B)** Wood necrosis area in wounded (W) and infected plants with *Neofusicoccum parvum* Bt-67 (NP) after one-month infection. **C)** Lignin content after one-month infection at the inoculation site at wounded and infected canes. **D)** Monolignol pathway showing the positions of cinnamic acid (CINN), *p*-coumaric acid (COUM), caffeic acid (CAFF), and ferulic acid (FER). **E)** Experimental design to screen for the surrender signal. Candidate monolignols are fed to *N. parvum* (NP) and the culture filtrate collected after 2 weeks is added to *V. rupestris* GFP-TuB6 as recipient to screen for phytotoxicity **(c)** Mortality of the recipient cells scored in response to culture filtrate from *N. parvum* after feeding the fungal donor with different monolignols as compared to culture filtrate from mock-treated cells (NP), or without any filtrate (CTRL).**F)** Mortality of the recipient cells in response to fungal metabolites extracted from mycelia (MYC) or recovered from the culture filtrates (CF) after fermentation for 24 hr with *p-*coumaric acid or ferulic acid to probe for potential differences in the secretion of the phytotoxins. Data represent means and SE from three independent biological experiments. Asterisks indicate statistical differences by LSD test at significance level with *P*<0.05 (*), *P*<0.01 (**). and *P*<0.001 (***) in **B** and **C**. Different letters represent statistical differences based on Duncan’s test with significant levels *P*<0.05 in **E** and **F**.

This was linked with lignin accumulation. Although drought induced lignin slightly (**Fig. 2b**, *W*), lignin content after infection (**Fig. 2b**, *NP*) rose further, slightly, but significantly. However, infection and drought act additively, not synergistically, with respect to lignin deposition.

### In response to ferulic acid, *N. parvum* secretes a FCA derivative

Since *N. parvum* aggressiveness depends on plant factors associated to the stress level of the plant, we tested different monolignol precursors for their ability to elicit the release of fungal phytotoxins (**Fig. 2d, e**). When we fermented the mycelia in presence of cinnamic acid, and added the culture-filtrate to *Vitis* cells, these cells displayed a mortality that was significantly lower than the mortality in response to filtrate from untreated mycelium. Likewise, both, *p-* coumaric acid and caffeic acid, reduced the toxicity of the fungal culture-filtrate down to the residual mortality of untreated cells (**Fig. 2d**), irrespective of precursor concentration (**Fig. S2**). However, ferulic acid, even at only 0.5 mM, boosted the toxicity of the culture-filtrate (**Fig. 2e)**. These mycelia also switched to the sexual cycle as evident from excessive spore release when ferulic acid was present in the medium. To test, whether ferulic acid and its precursor coumaric acid differ with respect to accumulation or to secretion, we separated medium from mycelia. Mycelium extract caused similar mortality rates in the target cells, no matter, whether the mycelia grew untreated or in presence of *p*-coumaric acid or ferulic acid (**Fig. 2f**). By contrast, the culture-filtrates from the same cells induced a different mortality: Fermentation with *p*-coumaric acid produced less phytotoxic culture-filtrate than that from *N. parvum* alone, while fermentation with ferulic acid caused higher mortality than the control culture-filtrate of *N. parvum* (**Fig. 2f**). This indicates that *p*-coumaric acid and ferulic acid do not differ with respect to their effect on phytotoxin biosynthesis, but with respect to phytotoxin secretion.

To identify the compound responsible for the phytotoxicity, we separated the fungal metabolites based on their hydrophobicity. Hereby, specific peaks appeared only after ferulic treatment and increased (**Fig. 3a**) depending on hydrophobicity (% MeCN) lacking from the control culture-filtrate of *N. parvum* (**Fig. S3**). For instance, a vanillic acid-like compound (**Fig. 3a**, orange box) increased first with rising concentrations of MeCN but was absent in the most hydrophobic fraction (100% MeCN), while a mellein-like compound increased up to 50% MeCN, but disappeared from 75% MeCN (**Fig. 3a**, blue box). Only one peak (**Fig. 3a**, red box, retention time 4,79 min) was prominent, and increased progressively with increasing the hydrophobicity of the extraction, qualifying as bioactive candidate, because the phytotoxicity of the fungal metabolites triggered by ferulic acid increased with the hydrophobicity of the fraction (**Fig. 3b**), with a very strong correlation (R= 0.99). After plotting the induced mortality over the area for the peak at retention time 4.79 min (**Fig. 3a**, red box), we found a clear saturation curve with a high correlation of R=0.9 **(Fig. 3c)**.

**Fig. 3.**
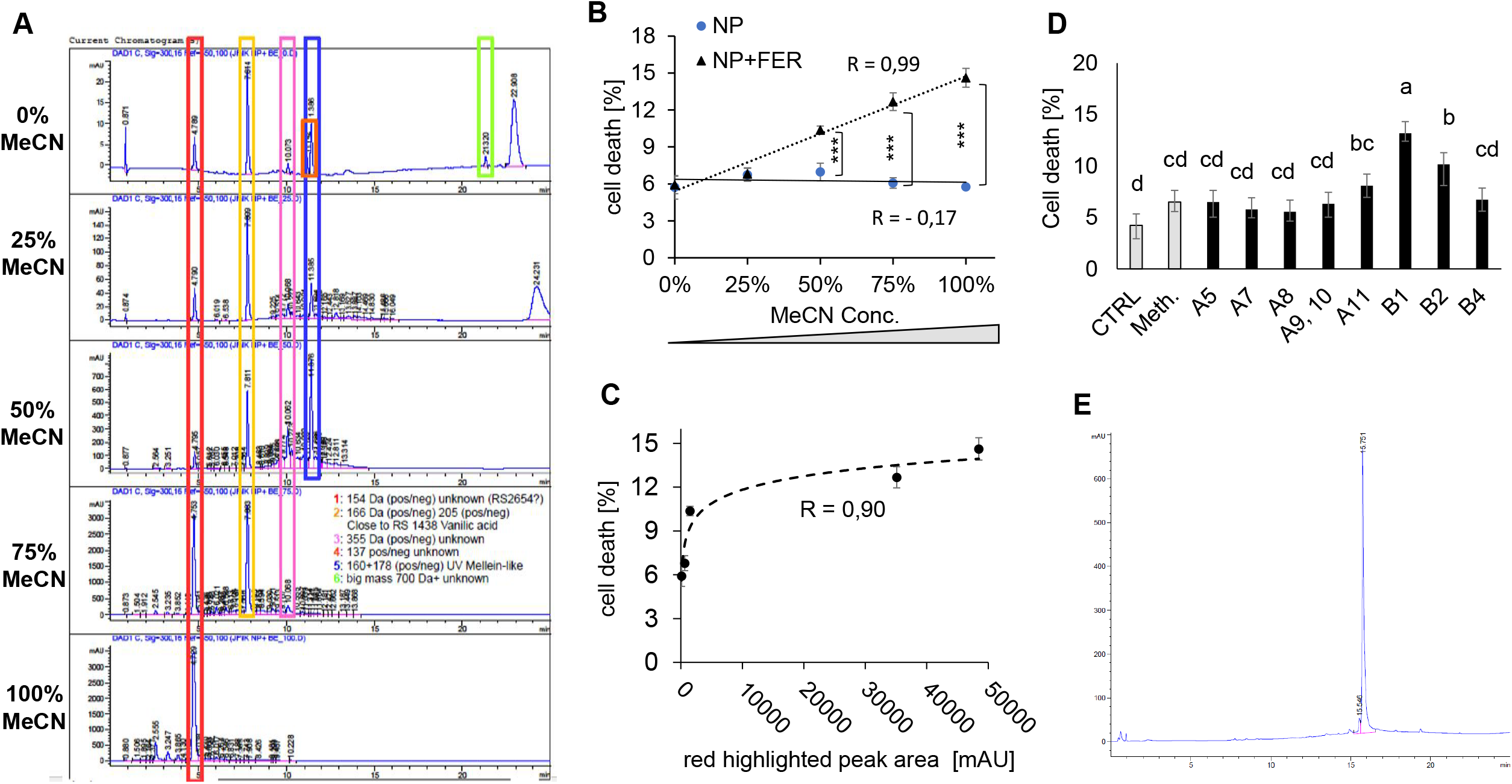
Identifying the fungal toxin released in response to fermentation with ferulic acid for 24 hr. **A)** HPLC UV readouts for the fungal metabolites secreted by *Neofusicocum parvum* Bt-67 in response to fermentation with 0.5 mM ferulic acid and fractionated using a solid phase extraction with a gradient of acetonitrile (0-100% MeCN). **B)** Mortality of *Vitis rupestris* GFP-TuB6 suspension cells in response to the fractions from fungal culture filtrates shown in **A**. For each fraction, a concentration of 20 µg/ml *Vitis* cells was administered. **C)** Correlation between mortality of the *Vitis* recipient cells and the area for the peak highlighted in **A** by the red box. **D)** Mortality of *V. rupestris* GFP-TuB6 cells in response to subfractions obtained from the most hydrophobic fraction of the acetonitrile gradient (MeCN 100%) of *N. parvum* fermented with ferulic acid. Here, a lower concentration (13.3 µg/ml Vitis cells was used, as compared to **B**, because the subfractions were more efficient and the material had to be safeguarded for structural elucidation. Mortality was scored after 24 h. **E)** Compound isolated from fractions B1 and B2 by LC-MS analysis and identified as a derivative of Fusicoccin A. Bars represent means and SE from three independent experimental series. Asterisks indicate statistical differences by LSD test at significance level with *P*<0.05 (*), *P*<0.01 (**). and *P*<0.001 (***) in **B**. Different letters represent statistical differences based on Duncan’s test with significant levels P<0.05 in **D**.

Consequently, the most hydrophobic phase was selected for further sub-fractionation by LC and characterisation by MS (absorption and mass spectra are given in **Fig. S4**). These fractions from LC were then further tested for their ability to induce mortality in the *Vitis* cells (**Fig. 2d**). Here, the highest bioactivity dwelled in fractions B1 and B2 (**Fig. 2d)**. HPLC-MS analysis identified the bioactive compound as a derivative of FCA (**Fig. 3e)**. In the following, we tried to elucidate the mode of action of FCA.

### FCA induces autolysis in grapevine cells

To get insight into the quality of mortality induced by FCA, we used double staining with Acridine Orange (AO, membrane permeable) and Ethidium Bromide (EB, membrane impermeable, DNA binding), classifying different stages of cell-death (**Fig. 4a**). In response to FCA, the frequency of cells in the 1^st^ stage, and dead cells (**Fig. 4f**) became more abundant in a dose-dependent manner. The frequency of cells in 2^nd^ stage was low and mostly constant through all time-points. Interestingly, this steady-state level was higher for 6 µM FCA (around 10-15%) comparing to the higher dose (around 5-10%). Together with the low incidence, this fact indicates that 2^nd^ stage is short-lived. A cell that is losing the tightness of the nuclear envelope against EB is doomed to timely death. When the response to FCA is speeding up, this will also reduce the steady-state level of 2^nd^ stage.

**Fig 4.**
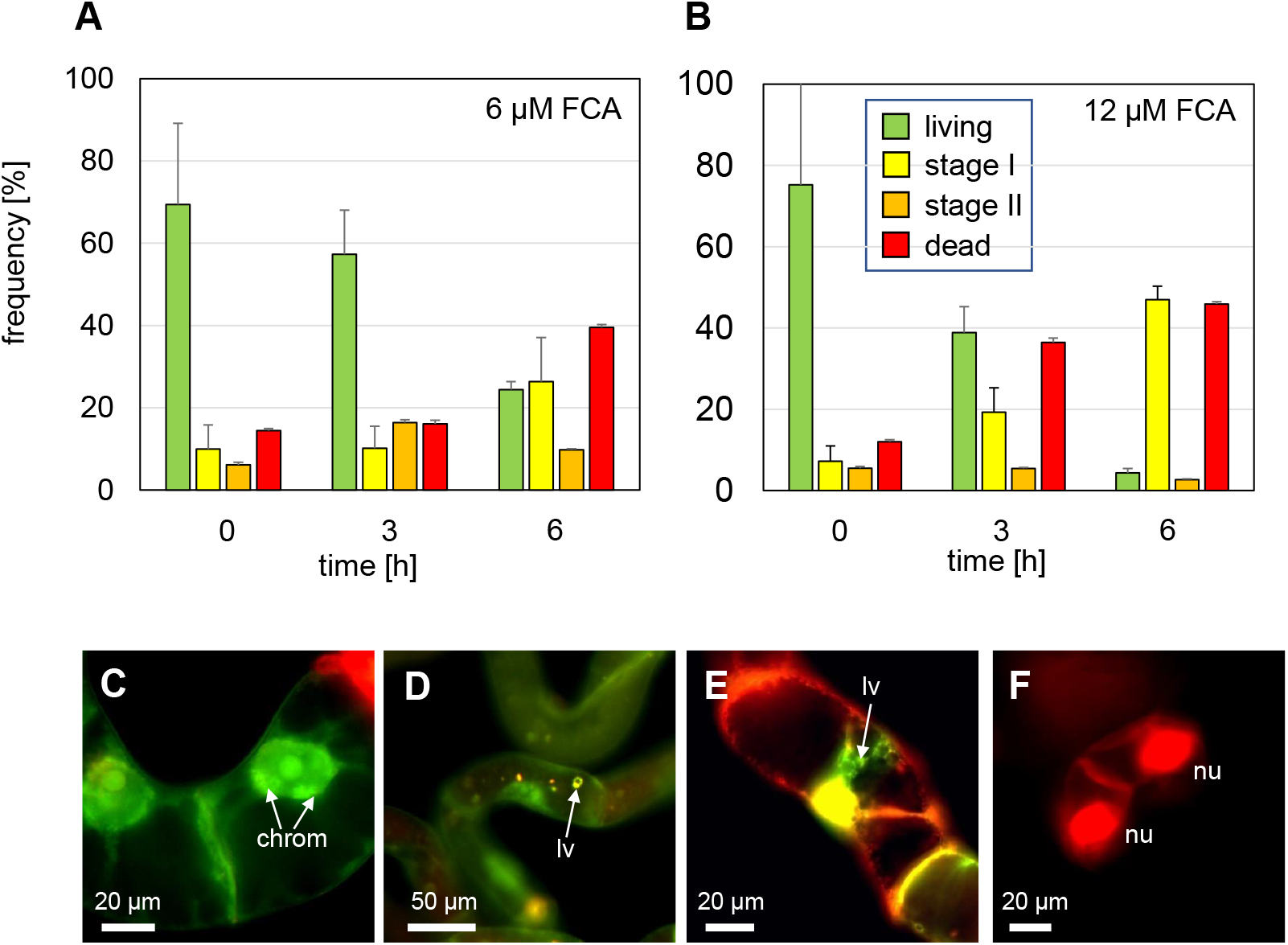
Cytological characterisation of the cell death in Vitis cells in response to Fusicoccin A. The cell death response was classified based on double staining with the membrane-permeable fluorochrome Acridine Orange (green signal) and the membrane-impermeable fluorochrome Ethidium Bromide (red signal). Living cells exclude Ethidium Bromide and appear green, cells in stage 1 show penetration of Ethidium Bromide into the cytoplasm, but still exclude the dye from the karyoplasm, cells in stage 2 show penetration of Ethidium Bromide into the nucleus, dead cells loose the Acridine Orange signal due to complete breakdown of the plasma membrane, while Ethidium Bromide remains sequestered at the DNA. Time course of these stages in response to 6 µM **(A)** and 12 µM **(B)** Fusicoccin A. Frequency distributions represent 300-400 individual cells collected from three independent experimental series. **(C-F)** progressive stages of autolytic cell death induced by Fusicoccin A, such as chromatin condensation in interphase nuclei (**C**, chrom), or appearance of lytic vacuoles (**lv**) in the cytoplasm (**D, E**). A dead cell void of cytoplasmic signals, but nuclei (**nu**) labelled by Ethidium Bromide, representing the terminal stage, is shown in **(F)**.

Moreover, this double staining assay allowed to observe transitional stages displaying cytological hallmarks of autolytic cell-death. For instance, transition to the 1^st^ stage was heralded by chromatin condensation (**Fig. 4c**), while progression through 1^st^ stage was accompanied by the formation of lytic vacuoles in the cytoplasm of dying cells (**Fig. 4d, e**). In the terminal stage, the nuclei were red, while the cytoplasmic signals observed in dying cells, vanished (**Fig. 4 c, d, e**).

### The response to FCA recapitulates cell-death related defence

We mapped the signalling deployed by FCA (6 µM), and observed a fast drop of extracellular pH, as it would be expected from activation of plasma-membrane (PM) proton ATPase. This acidification reached around -0.2 units within 30 min, kept a plateau for another 20 min and then underwent a second round of acidification reaching to -0.3 units at 90 min after addition of FCA (**Fig. 5a**). We further visualised superoxide by NBT staining and saw a rapid increase, significant already at 10 min and reaching to 25% within 1 h after FCA treatment (**Fig. 5b**), indicative of a stimulation of RBOH. After the peak at 60 min, the frequency of NBT positive cells dropped but remained elevated (three-fold of control cells). This drop might be linked with a loss of membrane integrity as found already substantial in the AO/EB staining (**Fig. 4a**).

**Fig 5.**
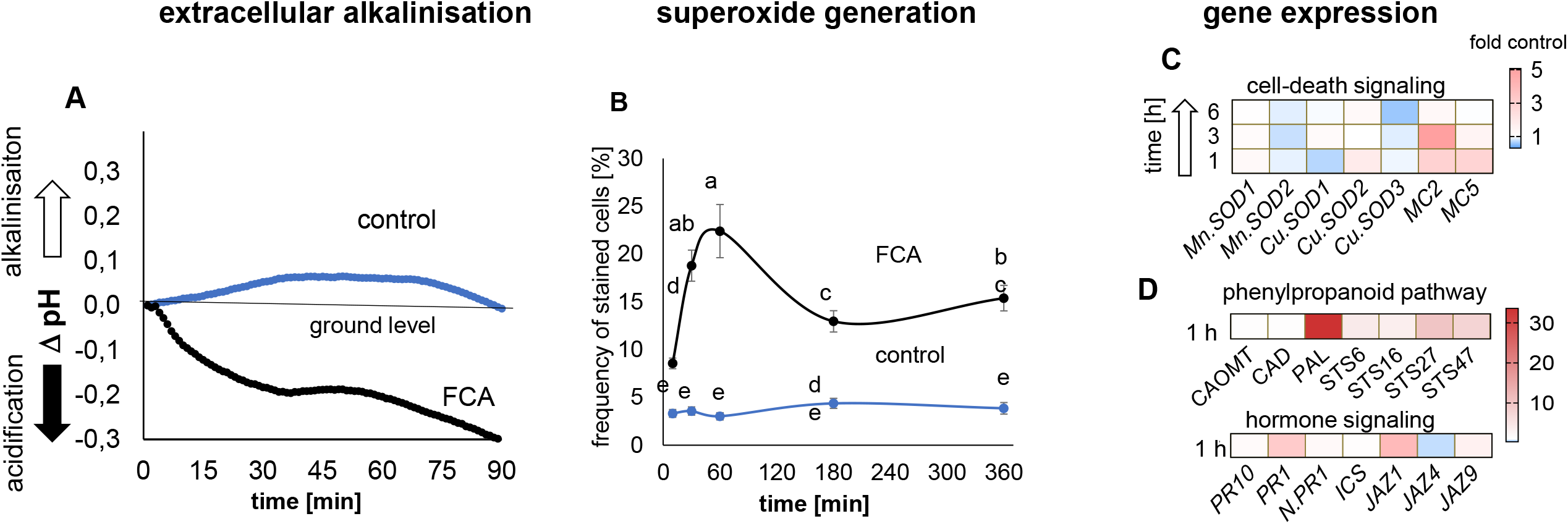
Rapid responses of *V. rupestris* GFP-TuB6 cells to Fusicoccin A (FCA). **A)** Activation of plasma-membrane proton ATPases by Fusicoccin A (6 µM) as detected by acidification of the extracellular medium. **B)** Detection of superoxide anions by 0.1% Nitroblue Tetrazolium. Data represent a population of 600-700 stained cells from three independent experimental series. **C)** Modulation of steady-state transcript levels of stress-marker genes measured by qPCR in response to Fusococcin A (6 µM) showing steady-state levels for transcripts involved in cell-death signalling such as mitochondrial (*MnSOD*) and plastidic (*CuSOD*) superoxide dismutase genes, as well as defence-related metacaspaces (*MC2, MC5*) over time. **D)** Transcripts of the phenyl propanoid pathway initiating from phenylammonium lyase (*PAL*), exemplarily probing monolignol synthesis (*CAOMT, CAD*) and stilbene synthesis (*STS6, STS16, STS27, STS47*). **E)** Transcripts of phytohormonal signaling genes (*JAZ1, JAZ2, JAZ9* for jasmonates; *PR1, ICS* for salicylic acid). Colour code represents the significant fold changes of 3 biological replicates normalised to the control of the respective time-point. Different letters represent statistical differences based on Duncan’s test with significant levels *P*<0.05.

We followed steady-state transcripts levels of potential stress-marker genes in response to FCA over time (**Fig. 5c**). Transcripts of *Superoxide Dismutase* (*MnSOD1* and *CuSOD2*) as markers for superoxide scavenging showed a rapid and steady induction, albeit to a mild extent. Also, *CuSOD1* displayed mild induction only transiently, at 3 h. The two metacaspases, *MC2* and *MC5*, markers for hypersensitive response (Gong *et al*., 2019) were rapidly (within 1 h) upregulated by FCA, but differed subsequently. While *MC5* declined later, the level of *MC2* transcripts continued to rise reaching the peak (5-fold) after 3 h.

The phenylpropanoid pathway gives rise to both, monolignols (i.e., the substrate of the fungus) and stilbenes (the central phytoalexin in grapevine). The transcripts of *PAL*, the first committed enzyme of the phenylpropanoid pathway, was massively (>30-fold) induced by FCA (**Fig. 5d**). Likewise, stilbene-biosynthesis transcripts accumulated significantly (10-20-fold) within 1 h, especially *STS27* and *STS47*. In contrast, the lignin biosynthesis genes, *CAOMT* and *CAD* did not show any notable induction.

In addition, FCA-challenged cells accumulated more transcripts of specific *JAZ* genes (*JAZ1* and *JAZ9*) (**Fig. 5d**). This indicates activation of jasmonate signalling as a hallmark of basal defence. Concerning salicylic acid, we observed a slight induction of *NPR1* and *PR10*, while *PR1* was clearly upregulated (~ 6-fold).

Thus, FCA induces a rapid extracellular acidification, and a rapid increase of superoxide, followed by induction of phytoalexin synthesis transcripts, and phytohormonal signalling (both, jasmonate and salicylic-acid), as well as metacaspases involved in hypersensitive cell-death. Therefore, FCA mimics several aspects of a cell-death response as normally observed in response to pathogen attack.

### FCA triggered PCD requires the activity of 14-3-3 proteins

The PM ATPases are activated by FCA through 14-3-3 proteins. Thus, blocking 14-3-3 proteins by the specific inhibitor BV02 (Stevers *et al*., 2018) should inhibit both acidification and PCD signalling. In fact, BV02 inhibited extracellular acidification in response to FCA (**Fig. 6a**) and modulated the responses of stress-marker genes (**Fig. 6b**). While the induction of *JAZ1* was not altered, both *STS27* and *STS47* were clearly amplified, while the induction of *PAL, PR1*, and the metacaspases *MC2* and *MC5* was mitigated. Furthermore, BV02 strongly reduced the mortality (**Fig. 6c**) from 30% (FCA alone) to 16% (FCA with BV02). Since the mortality induced by BV02 alone was around 20%, this low mortality meant that BV02 completely eliminated the mortality induced by FCA.

**Fig. 6.**
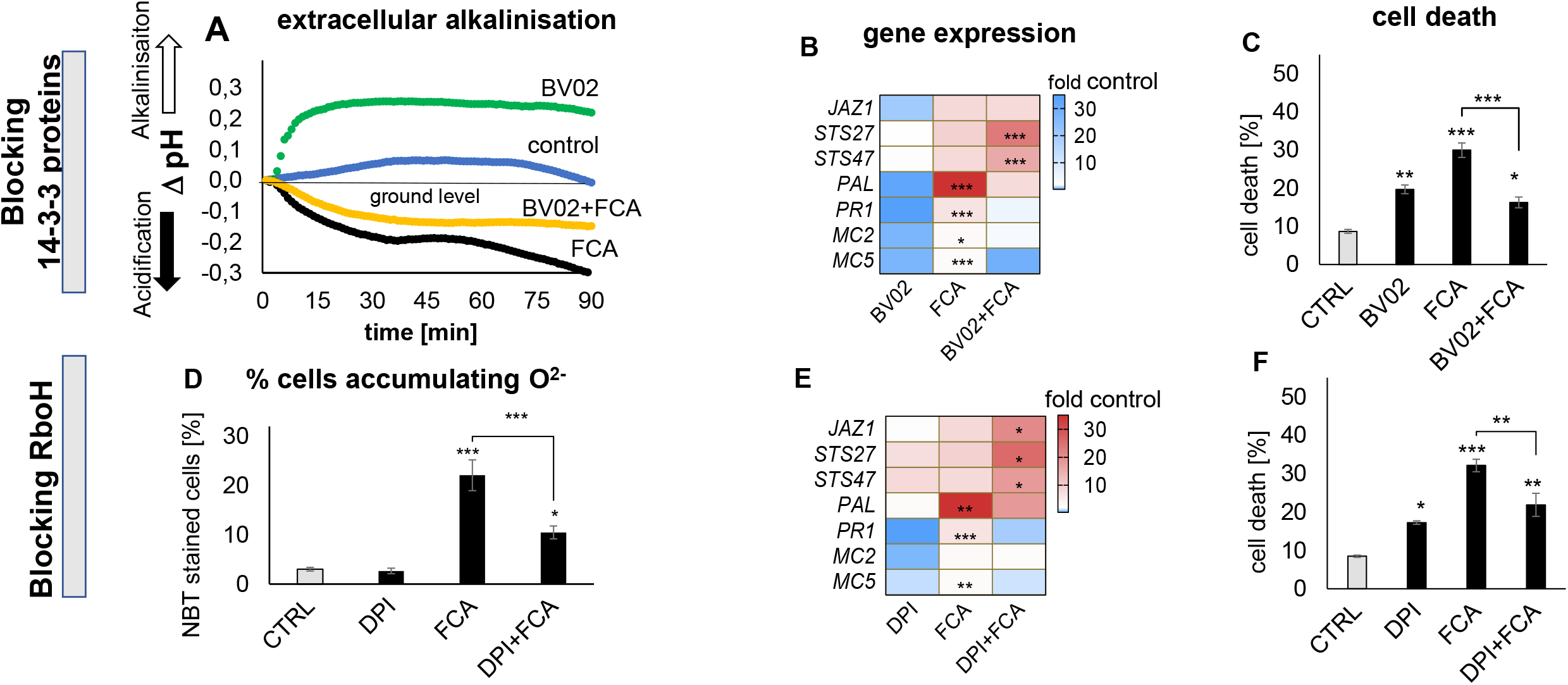
Role of 14-3-3 proteins and Respiratory burst oxygen Homologue for the cellular responses to Fusicoccin A. **A)** effect on extracellular acidification in response to 6 µM Fusicoccin A (FCA) as readout for plasma-membrane localised proton ATPases and after pre-treatment with 5 µM of the 14-3-3 inhibitor BV02 for 30 min. Effect of the 14-3-3 inhibitor BV02 **(B, C)** and the Respiratory burst oxidase homologue inhibitor diphenylene iodonium, DPI **(E, F)** on specific defence-related transcripts (**B, E**), and mortality (**C, F**). **D)** Effect of DPI on the generation of superoxide in response to FCA. BV02 was given in 5 µM, DPI in 1 µM, 60 min prior to adding FCA. Transcripts were scored after 1 h, mortality after 6 h. Color code represents means normalised to the control from 3 biological replicates, and asterisks indicate significant difference between Fusicoccin A alone versus the combination with the respective inhibitor. Bars represents means and SE from 3 biological replicates. Asterisks indicate statistical differences by LSD test at significance level with *P*<0.05 (*), *P*<0.01 (**). and *P*<0.001 (***).

### Blocking apoplastic oxidative burst modulates FCA signalling

Defence-related PCD is linked with apoplastic oxidative burst originating from RBOH, which should be disrupted by the RBOH inhibitor DPI. Superoxide production in response to FCA was reduced by DPI pre-treatment to ~ 50% (**Fig. 6d**), followed by specific effects on stress-marker genes (**Fig. 6e**). The basal defence genes *JAZ1, STS27*, and *STS47* were additively amplified by DPI and FCA. In contrast, DPI alone induced *PAL* transcripts by about 10-fold, but DPI reduced the induction by FCA to 50% of the level induced by FCA alone. A similar inhibition was observed for *PR1* and *MC5* transcripts, a mild (but insignificant) inhibition also for *MC2*. This second group (*PR1, MC2, MC5*) belonging to cell-death related defence (Chang and Nick, 2012; Gong *et al*., 2019), were not induced by DPI alone (contrasting with *JAZ1, STS27, STS47*, and *PAL* associated with basal immunity). Mitigation of the FCA responses of *PR1, MC2* and *MC5* by DPI should quell cell-death, which was the case (**Fig. 6f**).

### Microtubules respond to FCA and modulate FCA signalling

Since microtubules re-organise during defence and microtubule-directed compounds can modify defence responses (Chang and Nick, 2012), we studied the effect of FCA on microtubules in Vrup-TuB6 cells. FCA eliminated cortical microtubules within 30 min **(Fig. 7 a, d, g)**. This does not necessarily imply that microtubules participate in FCA signalling, because they might respond in a parallel pathway. To dissect this, we pre-stabilised microtubules first by taxol which significantly reduced microtubule elimination by FCA **(Fig. 6 d, e, g)** and saw also modulated FCA responses of defence-related transcripts (**Fig. 7h**). While taxol upregulated genes driving phytoalexin biosynthesis (*PAL, STS27* and *STS47*), and significantly amplified induction of these transcripts by FCA, the pattern for *MC2, MC5* and *PR1* transcripts differed qualitatively. Here, taxol significantly inhibited their induction by FCA. Although taxol stabilised microtubules, and modulated the gene expression in response to FCA, it could not mitigate the mortality driven by FCA **(Fig. 7i)**. However, comparing the mortality to FCA between our marker line *At*TUB6-GFP, to non-transformed *Vitis rupestris* cell lines, showed that ß-tubulin overexpression reduced mortality in response to FCA, to about 25% **(Fig. 7j)**. To test the effect of microtubules elimination on the mortality response to FCA, microtubules were eliminated by oryzalin within 30 min **(Fig. 7c)**. This elimination persisted during subsequent treatment with FCA **(Fig. 7f)** but did not change the mortality response to FCA (**Fig. S5**).

**Fig. 7.**
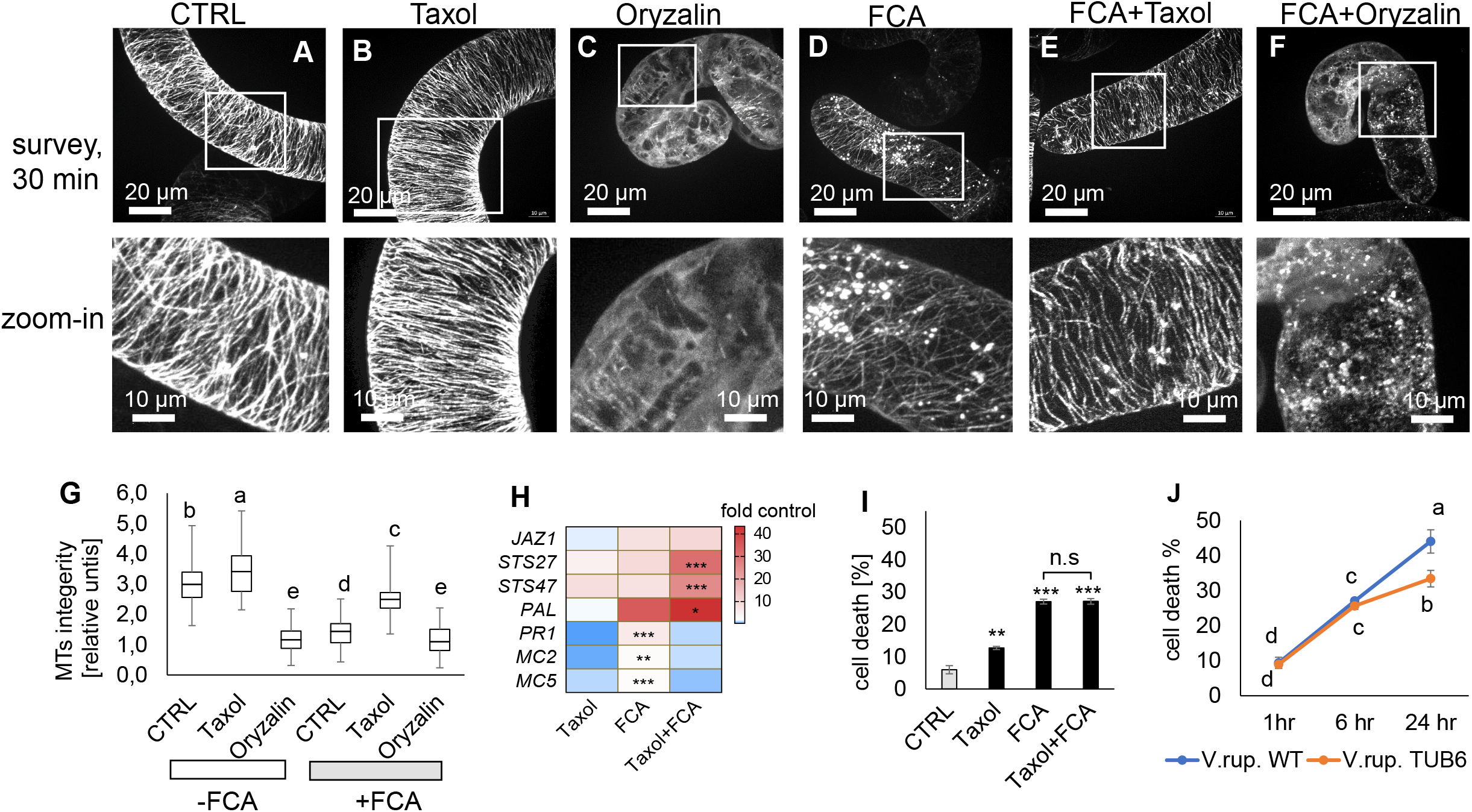
Microtubular response to Fusicoccin A, and role of microtubules for the cellular responses to Fusicoccin A. Representative *V. rupestris* GFP-TuB6 cells after 30 min of treatment with microtubule-modulating compounds either in the absence **(A-C)** or in presence of FCA **(D-F)**. Solvent controls **(A, D)** 0.1% DMSO, 10 µM taxol **(B, E)**, or 10 µM Oryzalin **(C, F)** are shown. **G)** quantification of microtubule integrity for the treatments shown representatively in **(A-F)**. Data represent medians, inter-quartiles, and extreme values for measurements from at least 30 individual cells. **H)** heat map of steady-state transcripts levels of stress-marker genes 1 h after addition of 6 µM FCA either alone or in combination with 10 µM Taxol. **I)** Mortality scored at 6 h after addition of either 6 µM FCA, 10 µM Taxol, or a combination of both compounds. J) Overexpression of microtubules in the *Vitis rupestris* cell line *At*TUB6-GFP led to less cell death rates due Fusicoccin A treatment comparing to *Vitis rupestris* wild type. Data represent mean and standard errors from three independent biological experiments comprising 1500 individual cells. Different letters represent statistical differences based on Duncan’s test with significant levels *P*<0.05 in **G** and **J**. Asterisks indicate statistical differences by LSD test at significance level with *P*<0.05 (*), *P*<0.01 (**). and *P*<0.001 (***) in **H** and **I**.

### Actin filaments respond to and modulate FCA signalling

Since actin filaments are implicated in PCD signalling (for review see Smertenko & Franklin-Tong, 2011; Chang *et al*., 2015), we used a grapevine actin-marker cell line to visualise actin filaments responses to FCA. While in control cells (**Fig. 8a**), actin was organised in a subcortical network of fine strands, it had repartitioned from the cortical network to bundled transvacuolar cables 90 min after FCA addition (**Fig. 8c**). This bundling was significant as seen from measuring bundle width (**Fig. 8e**). To probe whether actin bundling is necessary for FCA-triggered PCD, we eliminated actin strands by Latrunculin B pre-treatment (**Fig. 8b)** suppressing both the formation of actin cables in response to FCA (**Fig. 8e**) and FCA-dependent mortality to around half the value seen without latrunculin B (**Fig. 8g)**. Moreover, the FCA responses of defence-related genes for Latrunculin B pretreatment (**Fig. 8f**) were similar, but not identical to those seen for taxol pre-treatment (**Fig. 7h**). While both, Latrunculin B and taxol amplified the induction of *STS27* and *STS47* by FCA and both suppressed the induction of *MC5*, there were differences: Latrunculin B amplified the FCA response of *JAZ1*, but reduced that of *PAL*. Instead, taxol amplified *PAL*. Likewise, taxol could clearly suppress the induction of *MC2* and *PR1* by FCA, which was not seen for Latrunculin B **(Fig. 8f)**. The amplification of *STS27* and *STS47* is of particular interest because the stabilisation of microtubules and the elimination of actin strands have the same effect, indicating antagonistic roles of the two components of the cytoskeleton in processing the signal deployed by FCA.

**Fig. 8.**
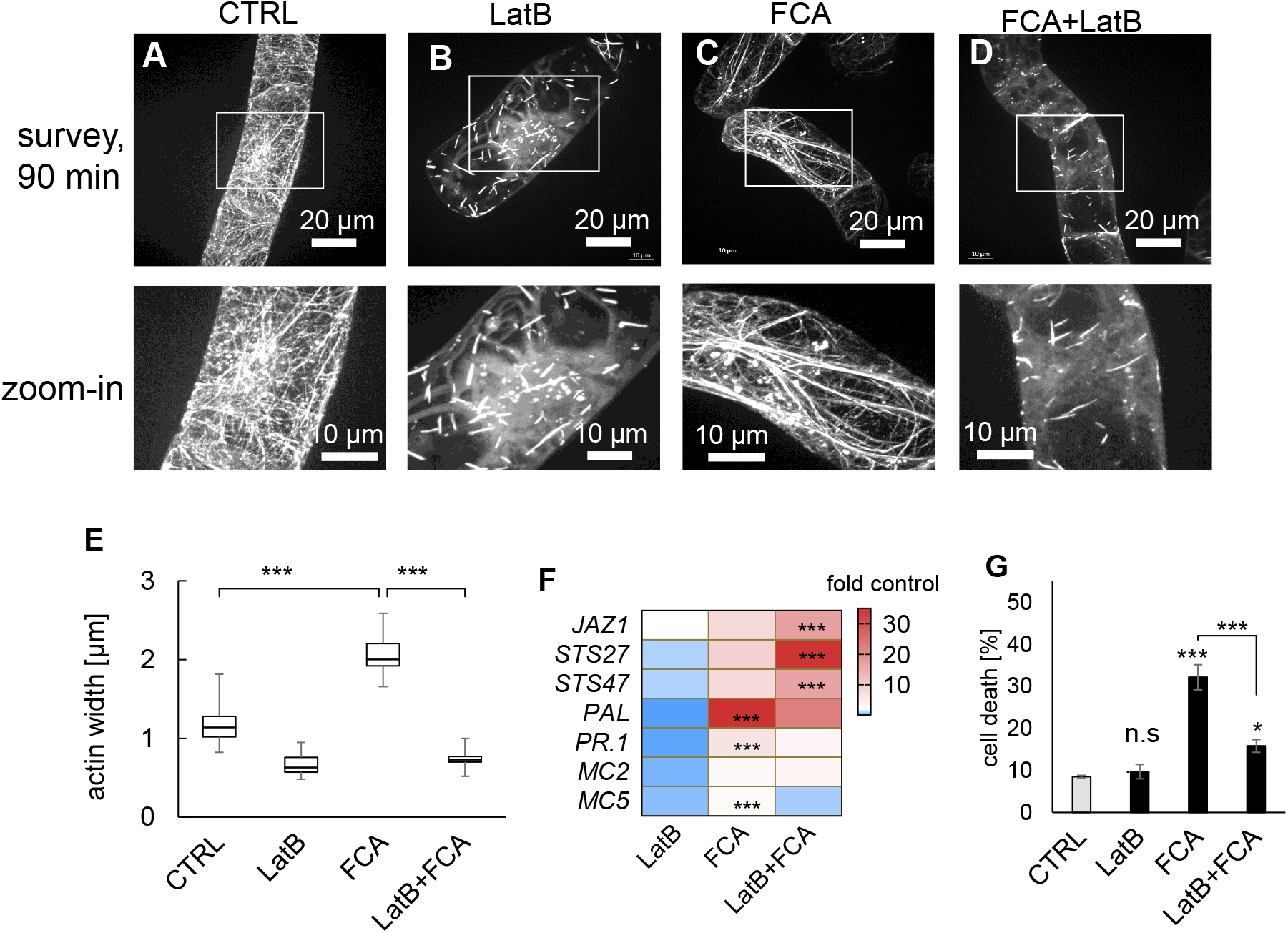
Actin response to Fusicoccin A, and role of actin for the cellular responses to Fusicoccin A (6 µM). Representative *V. vinifera* cv. Chardonnay FABD2-GFP cells after 90 min of treatment with the actin-eliminating compound Latrunculin B (2 µM) in the absence **(b)** or in presence of FCA **(d)**, compared to the solvent control **(a)** 0.1% DMSO, or to Fusicoccin A alone **(c). e)** quantification of actin bundling for the treatments shown representatively in **(a-d)**. Data represent medians, quartiles, and extreme values for measurements from at least 30 individual cells. **f)** heat map of steady-state transcripts levels of stress-marker genes 1 h after addition of 6 µM FCA either alone or in combination with 2 µM LatB. **i)** Mortality scored at 6 h after addition of either 6 µM FCA, 2 µM FCA, or a combination of both compounds. Data represent mean and standard errors from three independent biological experiments comprising 1500 individual cells. Asterisks indicate statistical differences based on LSD test with significant levels *P*<0.05 (*), *P*<0.01 (**), and *P*<0.001 (***).

### Overexpression of metacaspases increased cell-death triggered by FCA

In grapevine, defence-related PCD associates with the upregulation of two specific metacaspases, *MC2* and *MC5* (Gong *et al*., 2019). As FCA induced the transcript levels of these metacaspases genes, we tested whether these metacaspases participate in the cell-death driven by FCA. For this purpose, we used two BY-2 cell lines overexpressing these metacaspases from *V. rupestris*. These lines, MC2ox and MC5ox exhibited a significantly higher mortality in response to FCA, already manifest at the earliest tested time-point, 3 h (**Fig. 9a**). At 6 h, the mortality was tripled (MC2ox) or doubled (MC5ox) comparing to non-transformed WT, consistent with a causal role of these metacaspases in executing the cell-death response to FCA.

**Fig. 9.**
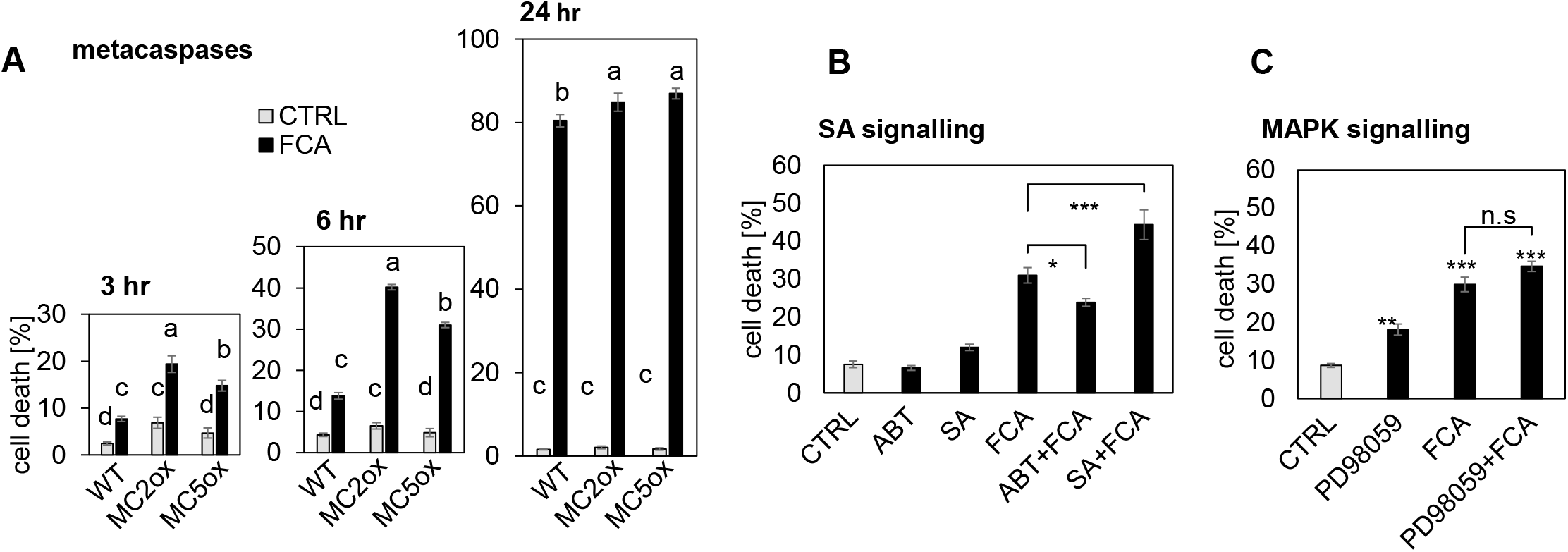
Probing for molecular components of the FCA response. **A)** Role of metacaspases. Time course of cell death in response to 6 µM Fusicoccin A in non transformed tobacco BY-2 cells (WT), and in cells overexpressing either metacaspase 2 (MC2ox) or metacaspase 5 (MC5ox) from *Vitis rupestris*. **B)** Role of Salicylic acid (50 µM) and its inhibitor, 1-aminobenzotriazole scored 6 h after addition of 6 µM Fusicoccin A to *V. rupestris* GFP-TuB6 cells. **C)** Role of MAPK signalling. Cell death scored 6 h after addition of 6 µM Fusicoccin A to *Vitis rupestris* GFP-TuB6 cells after pre-treatment with 50 µM of the MAPK inhibitor PD98059. Data represent means and SE from 3 biological replicates comprising 1500 individual cells per data point. Different letters represent statistical differences based on Duncan’s test with significant levels P<0.05 **A**, asterisks indicate statistical differences based on LSD test with significant levels P<0.05 (*), P<0.01 (**), and P<0.001 (***) in **B** and **C**.

### Salicylic Acid mediates FCA triggered mortality, MAPK cascades seem to be dispensible

Salicylic acid (SA) is often implicated in hypersensitive responses. We tested, therefore, how SA interacts with FCA induced cell death. The pre-treatment of SA prior to addition of FCA significantly increased mortality (**Fig. 9c)**. Since SA enhanced mortality in response to FCA, we tested the effect of blocking SA synthesis by pre-treatment with 25 µM ABT, an inhibitor of cinnamic acid 4-hydroxylase (Leon *et al*., 1995). ABT mitigated the mortality induced by FCA, indicative that endogenous salicylic acid is involved in the transduction of the FCA effect (**Fig. 9b)**. In contrast to SA, manipulation of MAPK signalling by the specific inhibitor PD98059, blocking basal immunity in grapevine (Chang and Nick, 2012), while rising mortality by itself (**Fig. 9c**), left the mortality response to FCA unaltered.

## 4. Discussion

The current work was motivated by a working model, where the apoplectic breakdown due to Grapevine Trunk Disease is a conditional phenomenon where the fungus changes from an endophytic towards a necrotrophic lifestyle, which implies that the endophyte must perceive and respond to input from the host (**Fig. 1**). This input can be operationally defined as surrender signal (although, like often in phytopathology, the sender is not releasing this signal on purpose). Using a cell-based experimental system based on grapevine suspension cells and the virulent model *Neofusicoccum parvum* Bt-67, we identified the surrender signal as ferulic acid, a monolignol induced under drought stress (Griesser *et al*., 2015). Ferulic acid (in contrast to its precursor coumaric acid) triggers in the fungus the release of a phytotoxin, FCA. We could show that this fungal compound acts as a signal evoking PCD in grapevine cells. Thus, the apoplectic breakdown can be described as a chemical communication between host and endophyte that shifts out of order under climate-born stress. This model stimulates the following questions: 1. What renders ferulic acid so specific as surrender signal? 2. By what mechanism can ferulic acid trigger the fungal release of FCA? 3. What is the functional context of FCA-triggered PCD? 4. What does this model contribute to a potential therapy against apoplectic breakdown.

### Why is ferulic acid an efficient surrender signal?

The conceptual model used in this study is based upon chemical signalling (**Fig. 1**). The chain of events driving disease outbreak is promoted by drought stress (**Fig. 2a**), generating a compound used by the fungus to sense that its host is doomed to irreversible stress. Searching for this “surrender signal”, we focussed on the phenylpropanoid pathway for two reasons: It gives rise to stilbenes, the major grapevine phytoalexins, also to lignin, the major carbon source for the fungus, and commonly accumulates under drought stress (Tu *et al*., 2020). The apposition of the hydrophobic lignin helps to retain water transport in vascular tissue and is, thus, of adaptive nature. The partitioning of phenylpropanoids between stilbenes versus lignin decides on the outcome of plant-fungal interaction (Khattab *et al*., 2021). In fact, we could show that feeding a monolignols precursor, ferulic acid, induced the fungus to release phytotoxins (**Fig. 2d-f**). Interestingly, cinnamic and coumaric acid, situated upstream of ferulic in the pathway, were even downmodulating the innate toxicity of the fungal culture-filtrate (**Fig. 2e**). Since coumaric acid is also the branching point for stilbenes, a high steady-state level of coumaric acid would report efficient synthesis of stilbenes, and, thus, host vigour. The same holds true, less tightly, for cinnamic acid (in fact, cinnamic acid is silencing phytotoxin release as well, albeit less efficiently compared to coumaric acid). The first metabolite committed for monolignols, is caffeic acid (www.kegg.jp, search vvi, ferulate). Thus, an increase in the steady-state levels of caffeic acid would report a bottleneck in lignin synthesis and, qualify as readout for the fungus to detect a serious host crisis. Unexpectedly, this is not, what we observe: caffeic acid is quelling phytotoxicity almost as efficiently as its precursor, coumaric acid. The key to this enigma may be the enzyme 4-coumarate ligase (gene VIT_16s0039g02040, protein UniProt F6HEF8), which is very permissive and accepts any phenolic acid as substrate (cinnamic acid, coumaric acid, caffeic acid, ferulic acid, 5-hydroxy ferulic acid (F5H), and even the monolignol sinapic acid, www.kegg.jp) to confer a Coenzyme A residue for further metabolisation. Changes in activity of this enzyme would, therefore, result in a complete shut-down of the entire pathway and, thus, is not apt to act as lever to sense changes in stilbene versus lignin partitioning. However, there exists an enzyme, specifically recruiting ferulic acid for monolignol synthesis: the cytochrome P_450_ 84A1 enzyme ferulate-5-hydroxylase (in grapevine present in tandem as gene VIT_03s0038g00500, protein UniProt D7U5I5 and the almost identical gene VIT_03s0038g00550, protein UniProt F6I194). Ferulate-5-hydroxylase is the rate-limiting enzyme for monolignol synthesis and as such strongly regulated (Ruegger *et al*., 1999). The fact that this enzyme was a prominent candidate during transcriptomics in response to *Neofusicoccum parvum* or with *Plasmopara viticola* (Massonnet et al., 2017; Liu *et al*., 2019), indicates that this gene is subject to tight regulation. The rice homologue of F5H is strongly downregulated under drought (tenor.dna.affrc.go.jp, search Os06g0349700), especially in roots. Whether this holds true for grapevine as well, is unknown, but would represent a testable implication of the surrender signal model.

### By what mechanism can ferulic acid trigger the fungal release of FCA?

The highly potent phytotoxin, FCA, is synthetised by cyclisation of geranylgeranyl diphosphate (GGDP) by an unusual diterpene synthase harbouring a C-terminal prenyltransferase domain (Toyomasu *et al*., 2007). In fact, a homologue with ~ 60% similarity and all the features of this fusicoccadiene synthase is present *N. parvum* genome (UniProt ID R1H2L0). How ferulic acid can trigger the release of FCA is unknown. However, it is a potent activator of fungal laccases that help the fungus forage lignin as carbon source (Piscitelli *et al*., 2011). Ferulic acid itself is broken down by feruloyl esterase into vanillin, which is taken up through a specific transporter (Shimizu *et al*., 2005). Feruloyl esterase can process coumaric acid, albeit at 10-fold reduced affinity, and 100-fold reduced velocity (Faulds *et al*., 2005). Thus, the suppressive activity of coumaric acid on phytotoxin secretion might be caused by coumaric acid blocking the substrate pocket of feruloyl esterase. The actual inducer of phytotoxin secretion might therefore be vanillin. In fact, this is supported by our observation that vanillic acid, the oxidised derivative of vanillin, accumulates in the toxic fractions of fungal exudates (**Fig. 3A**). The transporter for the uptake of vanillin has to be transported to the tip of the growing hyphae through the *Spitzenkörper*, controlling key steps in fungal development, including the transition towards conidia formation (for review see Harris, 2009). The foraging of ferulic acid as food source would correlate with developmental switches controlling secretion, and, thus, couple conidia formation with release of FCA, initiating apoplectic breakdown.

### What is the functional context of FCA-triggered PCD?

Our data introduce a new “surrender signal” into models of plant-pathogen dialogue (**Fig. 10**). While healthy plants accumulate coumaric acid, a precursor of bioactive stilbenes, keeping the fungus silent as peaceful endophyte, under climate-born stress, (drought), the monolignol precursor ferulic acid accumulates, signalling to the fungus the ensuing surrender of the host. In response, the fungus will convert ferulic acid into vanillic acid (Shimizu *et al*., 2005), which might trigger the *Spitzenkörper* to initiate transition to sexual development and secretion of the highly potent toxin, FCA, to kill the host for releasing the spores outside infected trunks **(Fig. 10. ➀, ➁)**. After binding to 14-3-3 proteins **(Fig. 10, ➂)**, FCA triggers extracellular acidification via binding to PM ATPases (Kinoshita and Shimazaki, 2001, **Fig. 10, ➃**). This will activate RBOH (Majumdar & Kar, 2018) and cause actin remodelling (Wang et al., 2021) leading to activation of defence genes. Microtubules act as negative regulators of signalling, possibly by tethering metacaspases and/or 14-3-3 proteins (Pignocchi and Doonan, 2011)), similar to their role in self-incompatibility (Poulter et al., 2008). While actin remodelling accompanies PCD in many systems (Smertenko and Tong, 2011), it does not induce PCD *per se* (Wang et al., 2021). It might be the recruitment of 14-3-3 proteins for the PM ATPases that releases the metacaspases. In fact, overexpression of type-II metacaspases in poplar is reducing the abundance of 14-3-3 proteins, indicative of a functional interaction (Bollhöner *et al*., 2018). Activation of metacaspases might then trigger expression of metacaspases genes sustaining PCD execution. Several, partially non-intuitive, implications of this model have been tested and confirmed. A rapid oxidative burst, inhibition of the responses by DPI, but also by latrunculin (destabilising actin) and taxol (stabilisting microtubules), but also the reduced mortality in cells overexpressing GFP-tagged tubulin. The model can also explain observations by others, such as the inhibition of FCA induced PCD sycamore cells by DPI (Malerba *et al*., 2008).

**Fig. 10.**
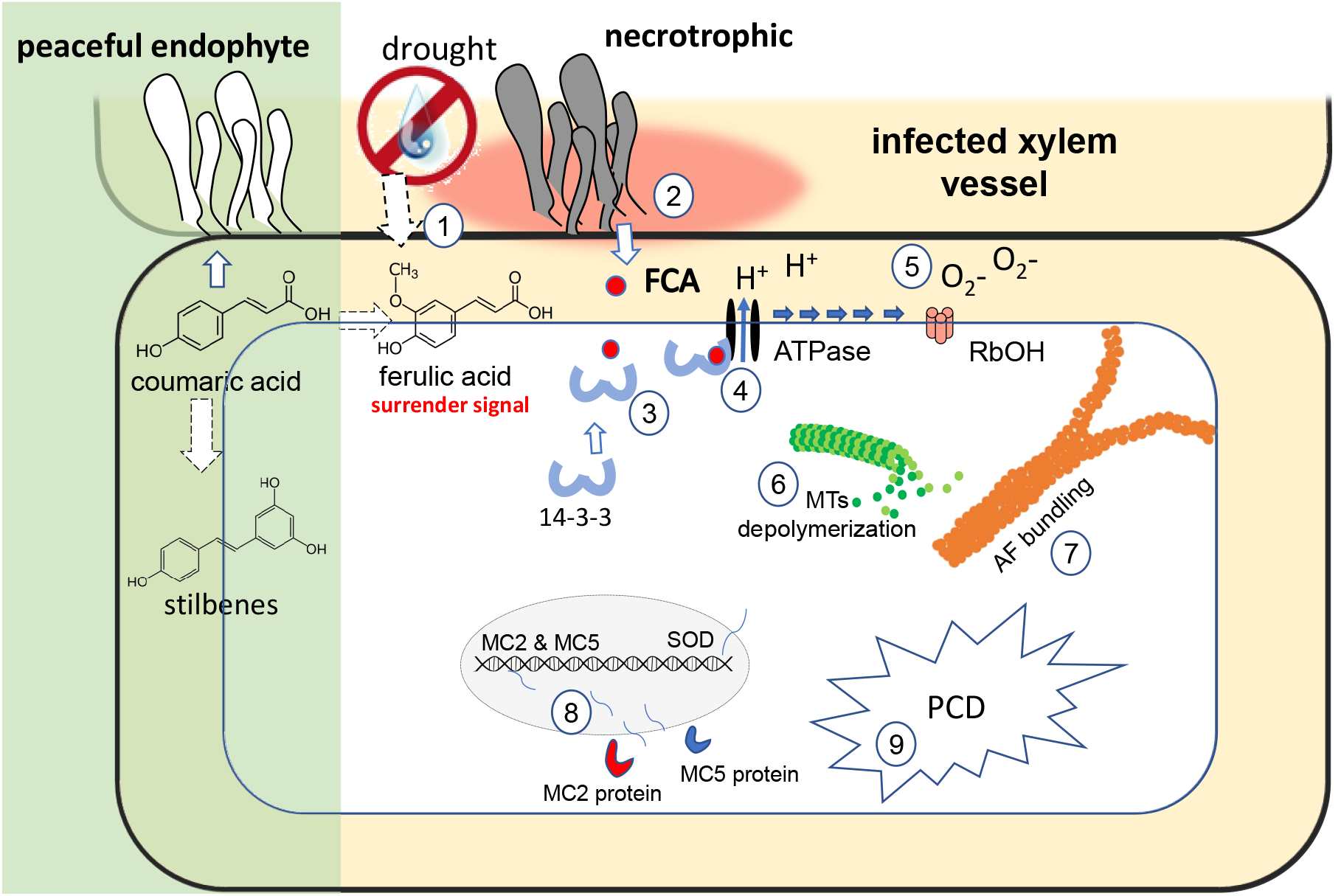
A visual model showing the chemical communication driving apoplexy in Botryosphaeriaceae-Vitis interaction and the stress signaling induced by the apoplexy signal (Fusicoccin A)

## 4. Outlook: What does this model contribute to a potential therapy against apoplectic breakdown

The outputs of this study pave the way for sustainable applications to control Grapevine Trunk Disease by channeling the phenylpropanoid pathway towards stilbene synthesis rather than lignification. To achieve this hypothesis-driven application, two approaches would be followed; (i) Chemical genetics as immediate strategy to contain disease outbreak to bridge the time until new resistant varieties become available. In this approach, compound libraries (van de Wouwer et al., 2016) will be used to modulate the pathway to avoid the accumulation of ferulic acid. (ii) Marker-assisted breeding based on genome sequencing of the almost entire population for the ancestral European Wild Grapevine (Liang et al., 2019), harbouring genes enhancing the synthesis of viniferins, which contain spread of GTDs (Khattab et al., 2021).

## Acknowledgements

This study was supported by the European Fund (Interreg Upper Rhine, projects Vitifutur, and DialogProTec). I.M.K. was awarded also a full PhD scholarship from the German Egyptian Research Long-term Scholarships DAAD-GERLS programme in addition to DAAD STIBET grant to complete this study.

## Supporting information

**Fig. S1**. The abundance of the fungal DNA in grapevines under drought stress

**Fig. S2**. Screening for the surrender signal among different concentration of the monolignols

**Fig. S3**. HPLC UV readouts for the fungal metabolites secreted by *Neofusicoccum parvum* Bt-67 in the absence of ferulic acid.

**Fig. S4**. Mass spectra for the toxic phase fractions of *Neofusicoccum parvum* Bt-67.

**Fig. S5**. Effect of Oryzalin on the cell mortality triggered by Fusicoccin A.

## Notes

### Competing Interest Statement

The authors have declared no competing interest.

## References

Akaberi S, Wang H, Claudel P, Riemann M, Hause B, Hugueney P, Nick P (2018) Grapevine Fatty Acid Hydroperoxide Lyase Generates Actin-Disrupting Volatiles and Promotes Defence-Related Cell Death. Journal of Experimental Botany, 69, 2883–2896.

Bollhöner B, Jokipii-Lukkari S, Bygdell J, Stael S, Adriasola M, Muñiz L, Breusegen F. V, Ezcurra I, Wingsle G, Tuominen H (2018) The function of two type II metacaspases in woody tissues of Populus trees. New Phytologist, 217(4), 1551–1565.

Bortolami G, Gambetta GA, Delzon S, Lamarque LJ, Pouzoulet J, Badel E, Burlett R, Charrier G, Cochard H, Dayer S, Jansen S (2019) Exploring the Hydraulic Failure Hypothesis of Esca Leaf Symptom Formation 1. Plant Physiology, 181(11), 1163–1174.

Buckel I, Andernach L, Schüffler A, Piepenbring M, Opatz T, Thines E (2017) Phytotoxic dioxolanones are potential virulence factors in the infection process of Guignardia bidwellii. Scientific Reports, 7(1), 8926.

Byczkowska A, Kunikowska A, Kaźmierczak A (2013) Determination of ACC-induced cell-programmed death in roots of Vicia faba ssp. minor seedlings by acridine orange and ethidium bromide staining. Protoplasma, 250(1), 121–128.

Carlucci A, Cibelli F, Lops F, Raimondo ML (2015) Characterization of Botryosphaeriaceae Species as Causal Agents of Trunk Diseases on Grapevines. Plant Disease, 99(12), 1678–1688.

Chang X, Heene E, Qiao F, Nick P (2011) The phytoalexin resveratrol regulates the initiation of hypersensitive cell death in vitis cell. PLoS ONE, 6(10).

Chang X, Nick P (2012) Defence signalling triggered by Flg22 and Harpin is integrated into a different stilbene output in Vitis cells. PLoS ONE, 7(7).

Chang X, Riemann M, Liu Q, Nick P (2015) Actin as deathly switch? How auxin can suppress cell-death related defence. PLoS ONE, 10(5), 1–22.

Djoukeng JD, Polli S, Larignon P, Abou-Mansour E (2009) Identification of phytotoxins from Botryosphaeria obtusa, a pathogen of black dead arm disease of grapevine. European Journal of Plant Pathology, 124(2), 303–308.

Duan D, Fischer S, Merz P, Bogs J, Riemann M, Nick P (2016) An ancestral allele of grapevine transcription factor MYB14 promotes plant defence. Journal of Experimental Botany, 67(6), 1795–1804.

Faulds CB, Molina R, Gonzalez R, Husband F, Juge N, Sanz-Aparicio J, Hermoso JA. (2005) Probing the determinants of substrate specificity of a feruloyl esterase, AnFaeA, from Aspergillus niger. FEBS Journal, 272(17), 4362–4371.

Gaff D, O. Okong’o-Ogola (1971) The use of non-permeating pigments for testing the survival of cells. Journal of Experimental Botany. 22:756–758.

Galarneau ERA, Lawrence DP, Travadon R, Baumgartner K (2019) Drought exacerbates botryosphaeria dieback symptoms in grapevines and confounds host-based molecular markers of infection by neofusicoccum parvum. Plant Disease, 103(7), 1738– 1745.

Gong P, Riemann M, Dong D, Stoeffler N, Gross B, Markel A, Nick P (2019) Two grapevine metacaspase genes mediate ETI-like cell death in grapevine defence against infection of Plasmopara viticola. Protoplasma, 256(4), 951–969.

Griesser M, Weingart G, Schoedl-Hummel K, Neumann N, Becker M, Varmuza K, Liebner F, Schuhmacher R, Forneck A (2015) Severe drought stress is affecting selected primary metabolites, polyphenols, and volatile metabolites in grapevine leaves (Vitis vinifera cv. Pinot noir). Plant Physiology and Biochemistry, 88, 17–26.

Guan P, Terigele Schmidt F, Riemann M, Fischer J, Thines E, Nick P (2020) Hunting modulators of plant defence: the grapevine trunk disease fungus Eutypa lata secretes an amplifier for plant basal immunity. Journal of Experimental Botany, (March).

Guan X, Buchholz G, Nick P (2015) Tubulin marker line of grapevine suspension cells as a tool to follow early stress responses. Journal of Plant Physiology, 176, 118–128.

Guan X, Essakhi S, Laloue H, Nick P, Bertsch C, Chong J (2016) Mining new resources for grape resistance against Botryosphaeriaceae: A focus on Vitis vinifera subsp. sylvestris. Plant Pathology, 65(2), 273–284.

Gómez P, Báidez AG, Ortuño A, Del Río JA (2016) Grapevine xylem response to fungi involved in trunk diseases. Annals of Applied Biology, 169(1), 116–124.

Harris SD (2009) The Spitzenkörper: A signalling hub for the control of fungal development? Molecular Microbiology, 73(5), 733–736.

Hofstetter V, Buyck B, Croll D, Viret O, Couloux A, Gindro K (2012) What if esca disease of grapevine were not a fungal disease? Fungal Diversity, 54(May), 51–67.

Jürges G, Kassemeyer HH, Dürrenberger M, Düggelin M, Nick P (2009) The mode of interaction between Vitis and Plasmopara viticola Berk. & Curt. Ex de Bary depends on the host species. Plant Biology, 11(6), 886–898.

Khattab IM, Sahi VP, Baltenweck R, Maia-Grondard A, Hugueney P, Bieler E, Dürrenberger M, Riemann M, Nick P (2021) Ancestral chemotypes of cultivated grapevine with resistance to Botryosphaeriaceae-related dieback allocate metabolism towards bioactive stilbenes. New Phytologist, 229(2), 1133–1146.

Kinoshita T, Shimazaki KI (2001) Analysis of the phosphorylation level in guard-cell plasma membrane H+-ATPase in response to fusicoccin. Plant and Cell Physiology, 42(4), 424–432.

Leon J, Shulaev V, Yalpani N, Lawton MA, Raskint I (1995) Benzoic acid 2-hydroxylase, a soluble oxygenase from tobacco, catalyzes salicylic acid biosynthesis (Nicotiana tabacum/tobacco mosaic virus/cytochrome P450/acquired resistance). Plant Biology, 92(October), 10413–10417.

Liang Z, Duan S, Sheng J, Zhu S, Ni X, Shao J, Liu C, Nick P, Du F, Fan P et al (2019) Whole-genome resequencing of 472 Vitis accessions for grapevine diversity and demographic history analyses. Nature Communications, 10(1), 1–12.

Lima MRM, Machado AF, Gubler WD (2017) Metabolomic Study of Chardonnay Grapevines Double Stressed with Esca-Associated Fungi and Drought. Phytopathology, 107(6), 669–680.

Liu R, Weng K, Dou M, Chen T, Yin X, Li Z, Li T, Zhang C, Xiang G, Liu G et al (2019) Transcriptomic analysis of Chinese wild Vitis pseudoreticulata in response to Plasmopara viticola. Protoplasma 256, 1409–1424.

Livak KJ, Schmittgen TD (2001) Analysis of relative gene expression data using real-time quantitative PCR and the 2(-Delta Delta C(T)) method. Methods 25, 402–408.

Loeffler F (1884) Untersuchung über die Bedeutung der Mikroorganismen für die Entstehung der Diphtherie beim Menschen, bei der Taube und beim Kalbe. Mittheilungen kaiserl Gesundheitsamt 2, 421–499.

Majumdar A, Kar RK (2018) Congruence between PM H+-ATPase and NADPH oxidase during root growth: a necessary probability. Protoplasma. Jul;255(4):1129–1137.

Malerba M, Contran N, Tonelli M, Crosti P, Cerana R (2008) Role of nitric oxide in actin depolymerization and programmed cell death induced by fusicoccin in sycamore (Acer pseudoplatanus) cultured cells. Physiologia Plantarum, 133(2), 449–457.

Massonnet M, Figueroa-Balderas R, Galarneau ERA, Miki S, Lawrence DP, Sun Q, Wallis CM, Baumgartner K, Cantu D (2017) Neofusicoccum parvum Colonization of the Grapevine Woody Stem Triggers Asynchronous Host Responses at the Site of Infection and in the Leaves. Frontiers in Plant Science, 8(June).

Mugnai L, Graniti A, Surico G (1999) Esca (Black Measles) and Brown Wood-Streaking: Two Old and Elusive Diseases of Grapevines. Plant Disease, 83(5), 404–418.

Pietrowska E, Różalska S, Kaźmierczak A, Nawrocka J, Małolepsza U (2014) Reactive oxygen and nitrogen (ROS and RNS) species generation and cell death in tomato suspension cultures—Botrytis cinerea interaction. Protoplasma, 252(1), 307–319.

Pignocchi C, Doonan JH (2011) Interaction of a 14-3-3 protein with the plant microtubule-associated protein EDE1. Annals of Botany, 107(7), 1103–1109.

Piscitelli A, Giardina P, Lettera V, Pezzella C, Sannia G, Faraco V (2011) Induction and transcriptional regulation of laccases in fungi. Curr Genomics. 12(2):104–112.

Poulter NS, Vatovec S, Franklin-Tong VE (2008) Microtubules are a target for self-incompatibility signaling in Papaver pollen. Plant Physiology, 146(3), 1358–1367.

Pouzoulet J, Scudiero E, Schiavon M, Rolshausen PE (2017) Xylem vessel diameter affects the compartmentalization of the vascular pathogen Phaeomoniella chlamydospora in grapevine. Frontiers in Plant Science, 8(August), 1–13.

Qiao F, Chang XL, Nick P (2010) The cytoskeleton enhances gene expression in the response to the Harpin elicitor in grapevine. Journal of Experimental Botany, 61(14), 4021–4031.

Ruegger M, Meyer K, Cusumano JC, Chapple C (1999) Regulation of Ferulate-5-Hydroxylase Expression in Arabidopsis in the Context of Sinapate Ester Biosynthesis, Plant Physiology, 119: 101–110.

Schwarzerová K, Zelenková S, Nick P, Opatrný Z (2002) Aluminum-induced rapid changes in the microtubular cytoskeleton of tobacco cell lines. Plant and Cell Physiology, 43(2), 207–216.

Shimizu M, Kobayashi Y, Tanaka H, Wariishi H (2005) Transportation mechanism for vanillin uptake through fungal plasma membrane. Applied Microbiology and Biotechnology, 68(5), 673–679.

Smertenko A, Franklin-Tong VE (2011) Organisation and regulation of the cytoskeleton in plant programmed cell death. Cell Death and Differentiation, 18(8), 1263–1270.

Stempien E, Goddard ML, Wilhelm K, Tarnus C, Bertsch C, Chong J (2017) Grapevine Botryosphaeria dieback fungi have specific aggressiveness factor repertory involved in wood decay and stilbene metabolization. PLoS ONE, 12(12), 1–22.

Slippers B, Wingfield MJ (2007) Botryosphaeriaceae as endophytes and latent pathogens of woody plants: diversity, ecology and impact. Fungal Biology Reviews, 21(2–3), 90–106.

Steffens B, Sauter M (2009) Epidermal cell death in rice is confined to cells with a distinct molecular identity and is mediated by ethylene and H2O2 through an autoamplified signal pathway. Plant Cell, 21(1), 184–196.

Stevers LM, Sijbesma E, Botta M, Mackintosh C, Obsil T, Landrieu I, Cau Y, Wilson AJ, Karawajczyk V, Eickhoff J et al (2018) Modulators of 14-3-3 Protein-Protein Interactions. Journal of Medicinal Chemistry, 61(9), 3755–3778.

Svyatyna, K., Jikumaru, Y., Brendel, R., Reichelt, M., Mithöfer, A., Takano, M., Kamiya Y, Nick P, Riemann, M (2014) Light induces jasmonate-isoleucine conjugation via OsJAR1-dependent and -independent pathways in rice. Plant, Cell and Environment, 37(4), 827–839.

Toyomasu T, Tsukahara M, Kaneko A, Niida R, Mitsuhashi W, Dairi T, Kato N, Sassa T (2007) Fusicoccins are biosynthesized by an unusual chimera diterpene synthase in fungi. Proceedings of the National Academy of Sciences of the United States of America, 104(9), 3084–3088.

Tu M, Wang X, Yin, W. Yin W, Wang Y, Li Y, Zhang G, Li Z, Song J, Wang X (2020) Grapevine VlbZIP30 improves drought resistance by directly activating VvNAC17 and promoting lignin biosynthesis through the regulation of three peroxidase genes. Horticultural Research, 7, 150.

Úrbez-Torres JR, Gubler WD (2009) Pathogenicity of Botryosphaeriaceae species isolated from grapevine cankers in California. Plant Disease, 93(6), 584–592.

Úrbez-Torres JR (2011) The status of Botryosphaeriaceae species infecting grapevines José. Phytopathologia Mediterranea, 50(4).

van de Wouwer D, Vanholme R, Decou R, Goeminne G, Audenaert D, Nguyen L, Höfer R, Pesquet E, Vanholme B, Boerjan W (2016) Chemical genetics uncovers novel inhibitors of lignification, including p-iodobenzoic acid targeting CINNAMATE-4-HYDROXYLASE. Plant Physiology, 172(1), 198–220.

Wang H, Riemann M, Liu Q, Siegrist J, Nick P (2021) Glycyrrhizin, the active compound of the TCM drug Gan Cao stimulates actin remodelling and defence in grapevine. Plant Science, 302(April 2020), 110712.

Wang L, Nick P (2017) Cold sensing in grapevine—which signals are upstream of the microtubular “thermometer.” Plant Cell and Environment, 40(11), 2844–2857.

Watanabe N, Lam E (2011) Arabidopsis metacaspase 2d is a positive mediator of cell death induced during biotic and abiotic stresses. Plant Journal, 66(6), 969–982.

