## Supplemental figures for "Hunting the plant surrender signal activating apoplexy in grapevines after *Neofusicoccum parvum* infection"

**The following supporting figures is available for this article:**

### Fig. S1

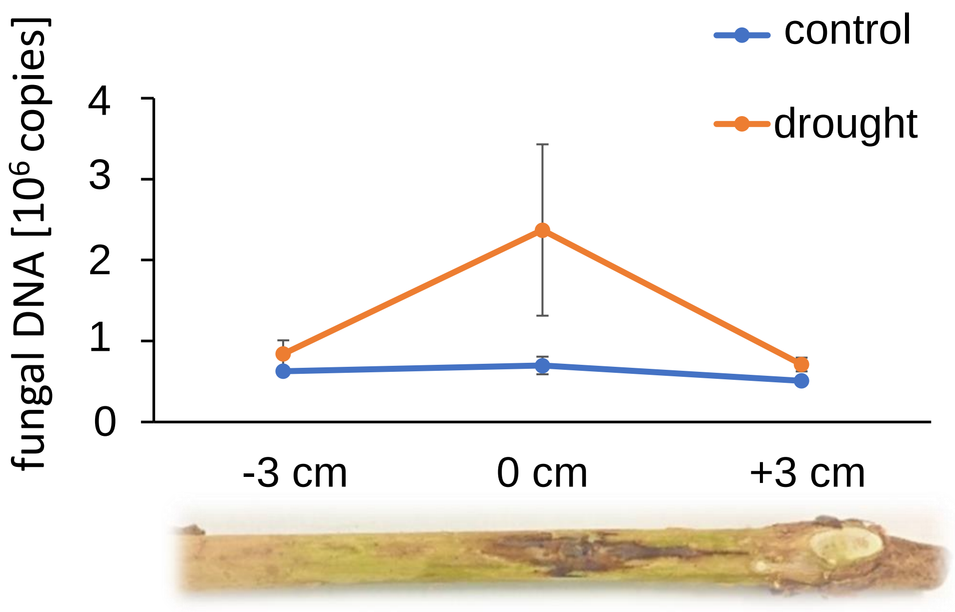


**Fig. S1.** Abundance of *Neofusicoccum parvum* Bt-67 in the site of inoculation (0 cm), or 3 cm below (-3 cm) or above (+3 cm) one month after infection in the host *vinifera* variety Augster Weiß either under standard irrigation (control) or drought stress (irrigation restrained to 20%). Data represent mean and standard errors from 3 individual plants.

### Fig. S2

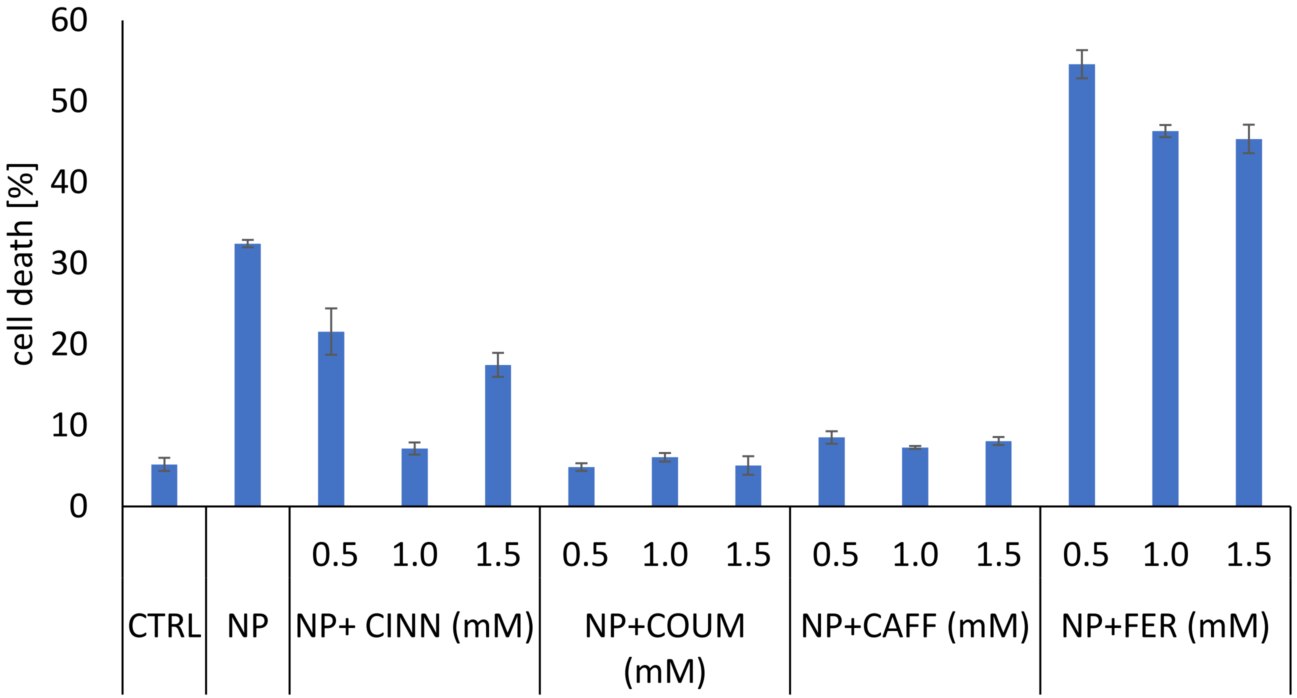


**Fig. S2.** Mortality in *V. rupestris* cells 24 h after addition of sterilised *Neofusicoccum parvum* Bt-67 culture filtrate (35 µl filtrate/ 1 ml *Vitis* cells) from mycelium fermented separately for 2 weeks with the indicated concentrations (0.5:1:1,5) mM of the four lignin precursors (cinnamic acid, *p*-coumaric acid, caffeic acid, ferulic acid). Bars represents means and SE from 3 biological replicates.

### Fig. S3

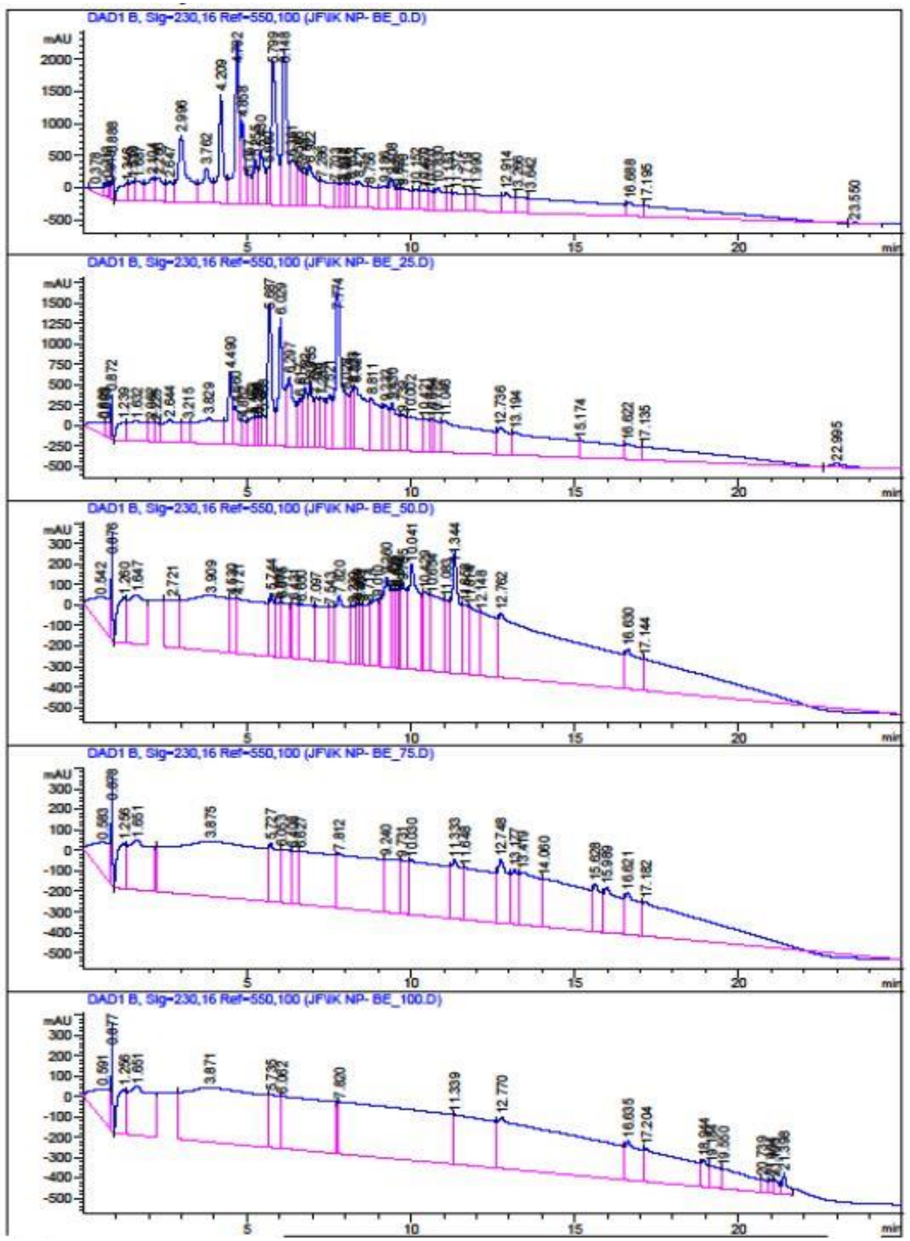


**Fig. S3.** HPLC UV readouts for the fungal metabolites secreted by *Neofusicoccum parvum* Bt-67 in the absence of ferulic acid and extracted using a solid extraction phase with a gradient of acetonitrile, MeCN (0%:25%:50%:75%:100%).

### Fig. S4.1

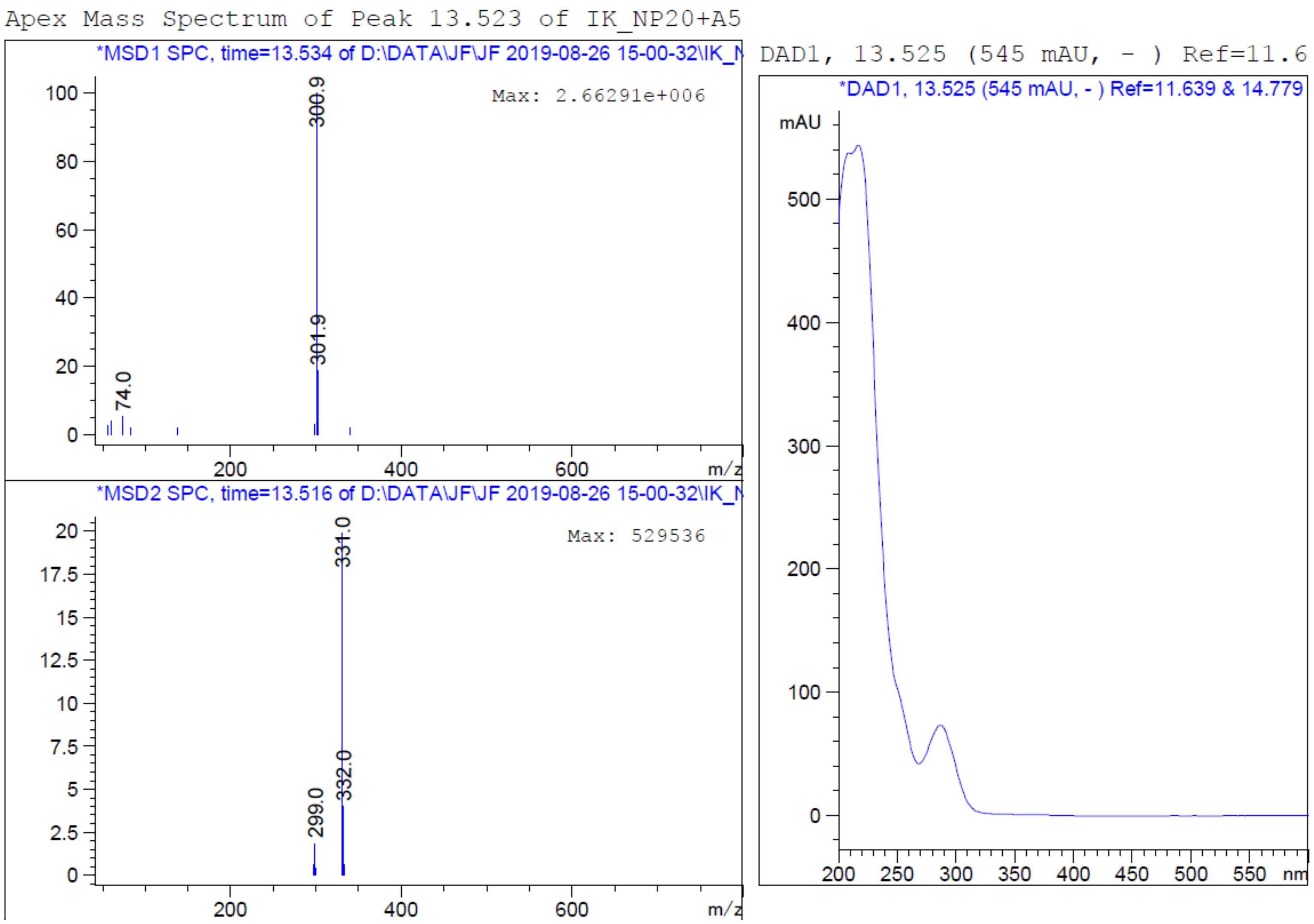


Mass spectra for fraction A5

#### Fig. S4.2

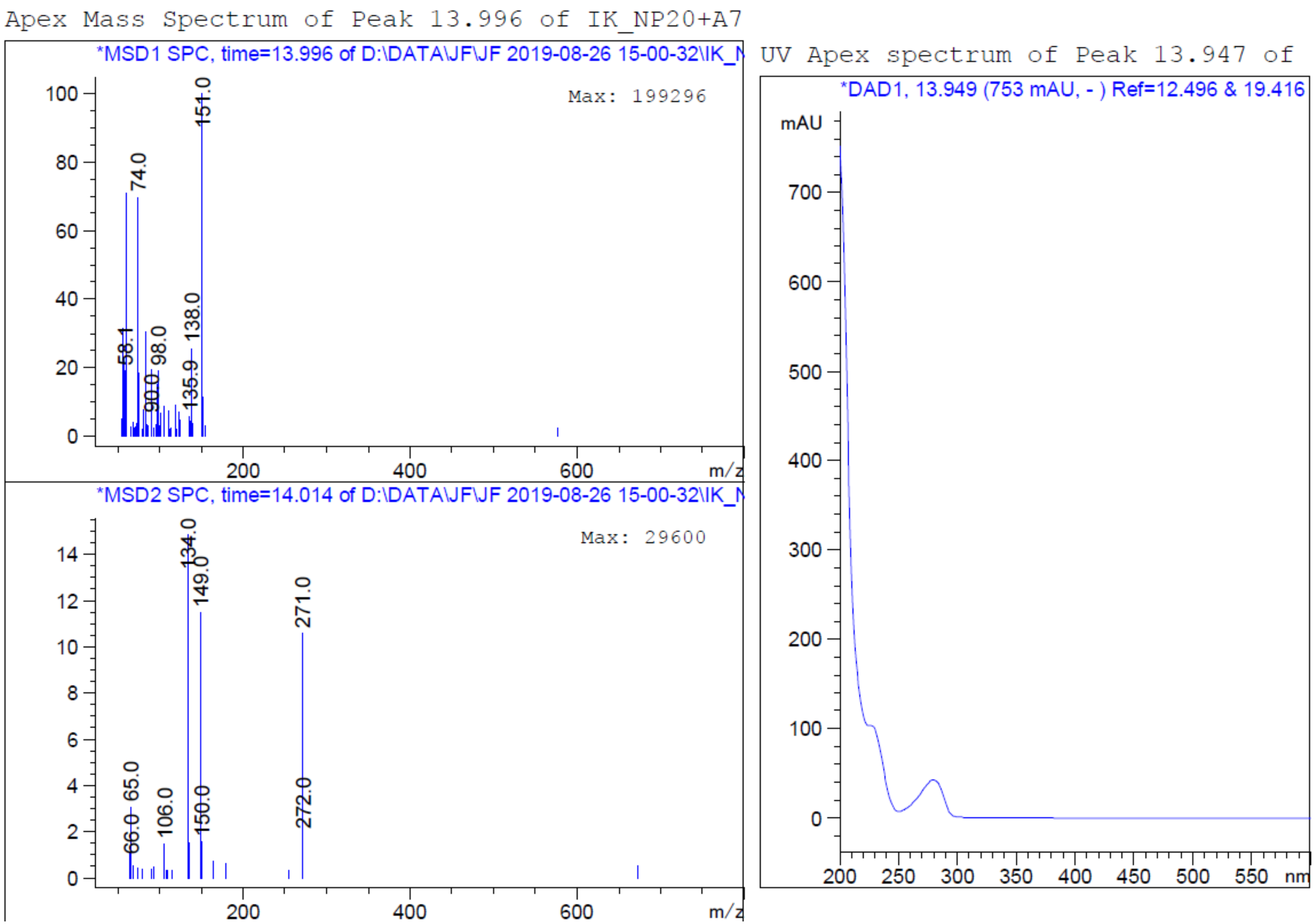


Mass spectra for fraction A7

### Fig. S4.3

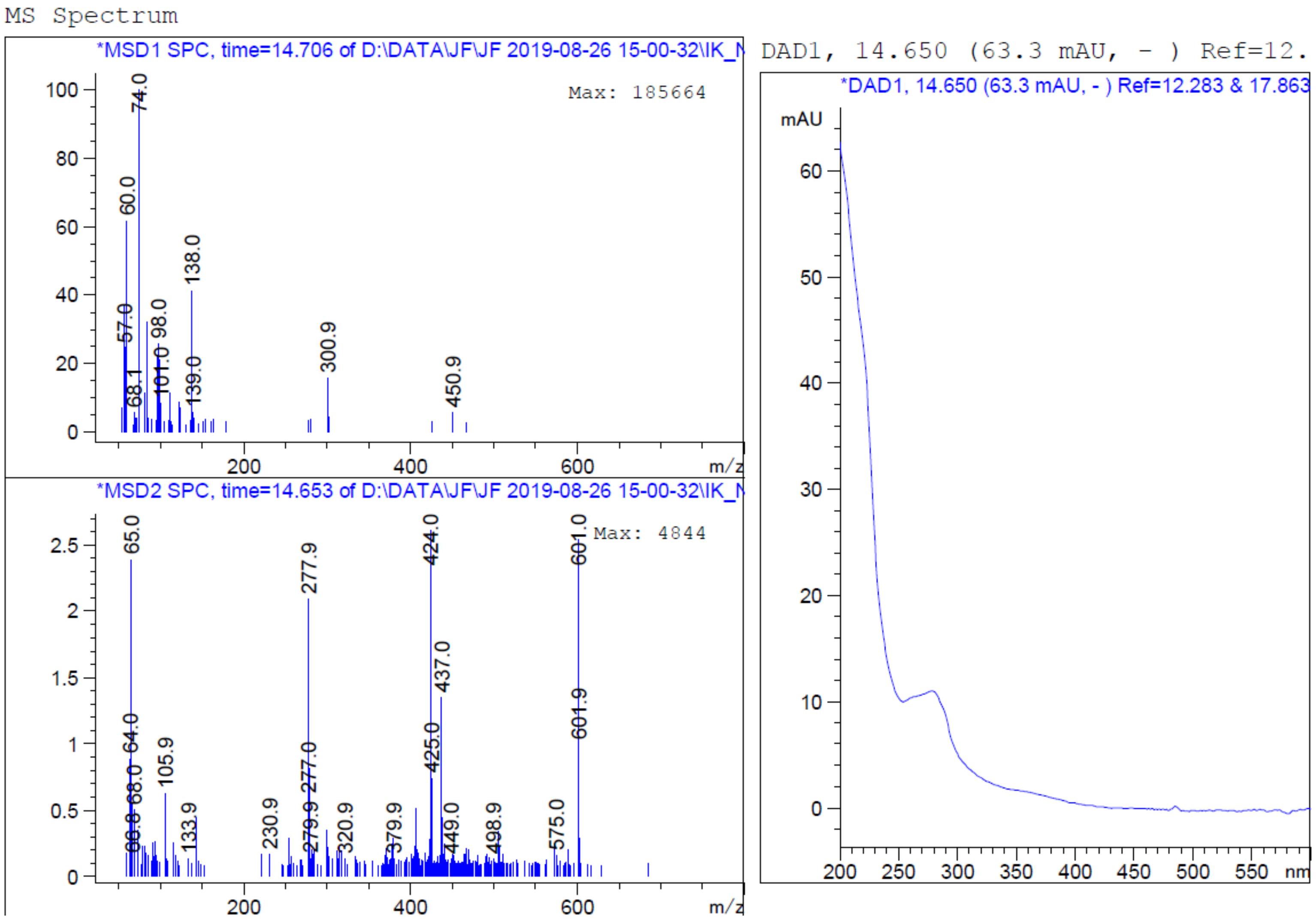


Mass spectra for fraction A8

### Fig. S4.4

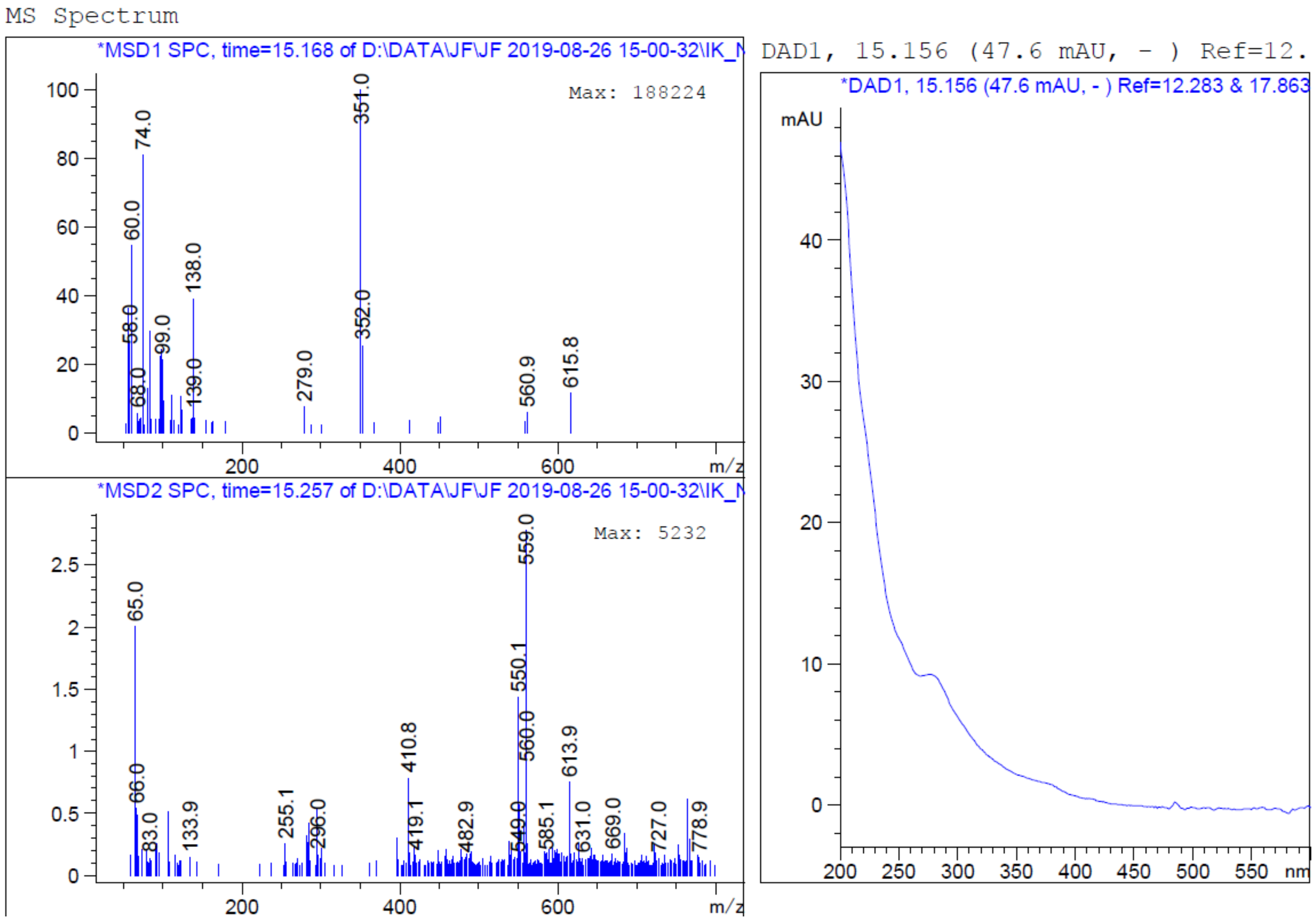


Mass spectra for fraction A8

### Fig. S4.5

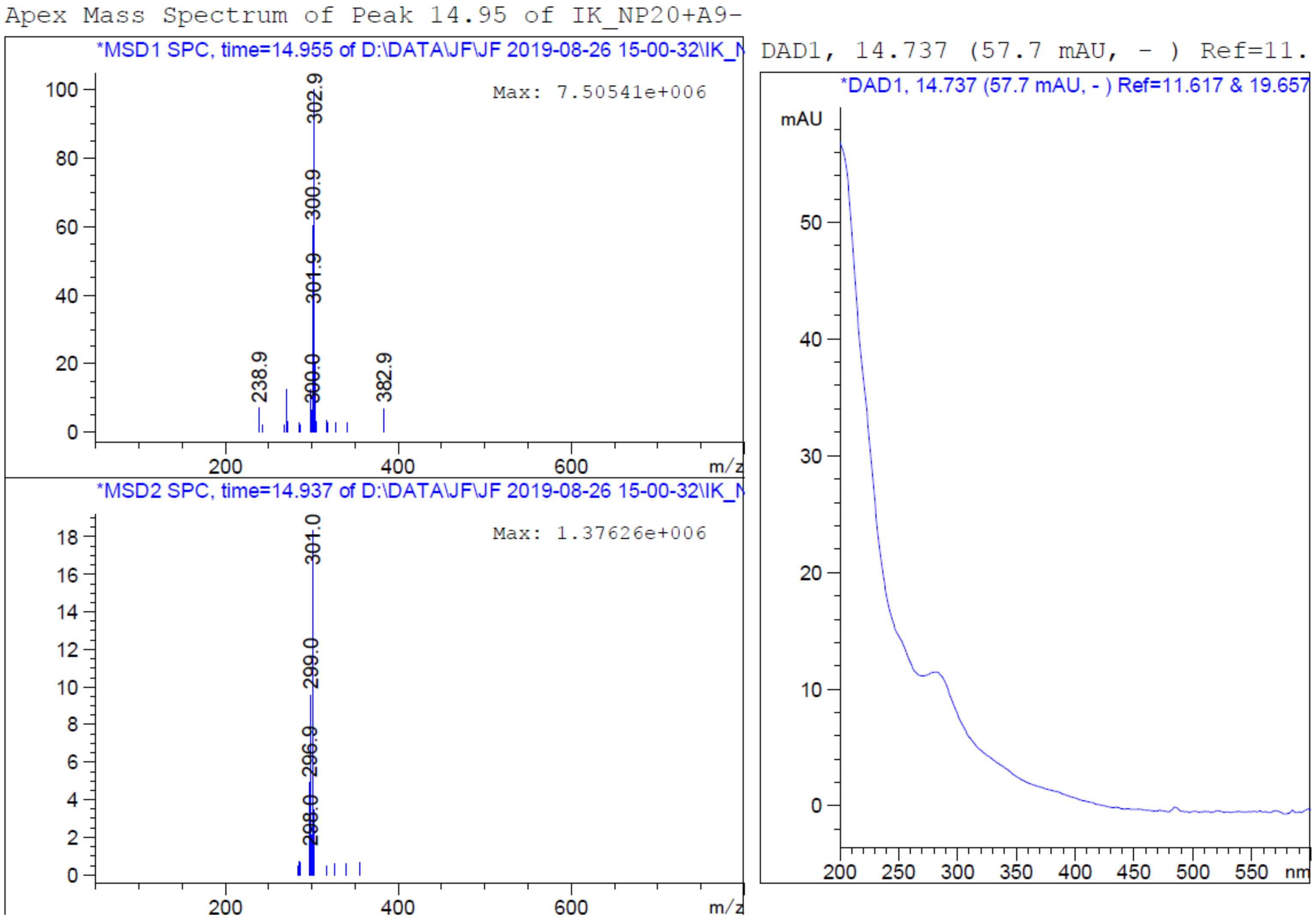


Mass spectra for fraction A9 & A10

### Fig. S4.6

The most toxic fractions, B1, B2

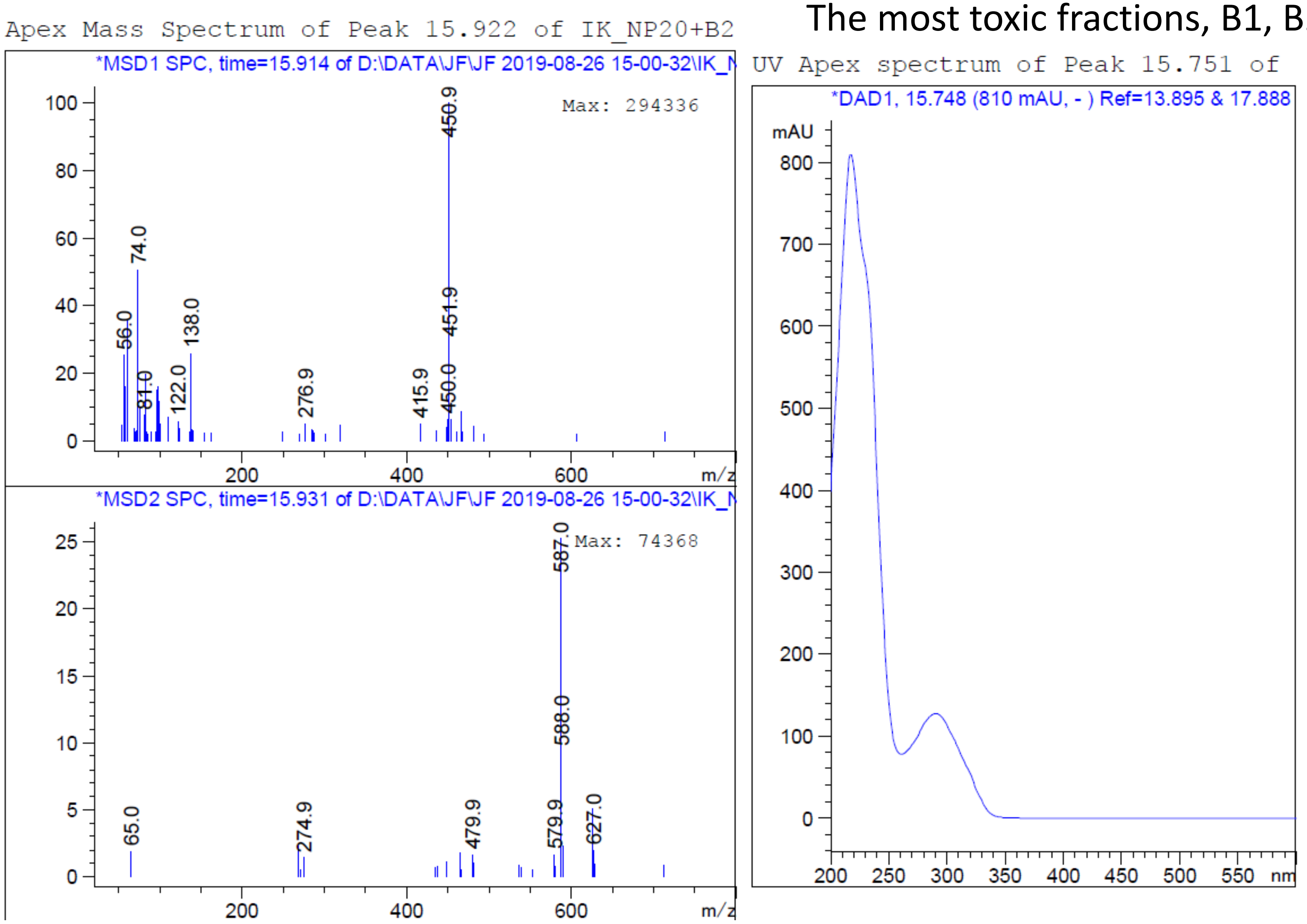


Mass spectra for fraction B1 & B2

#### Fig. S4.7

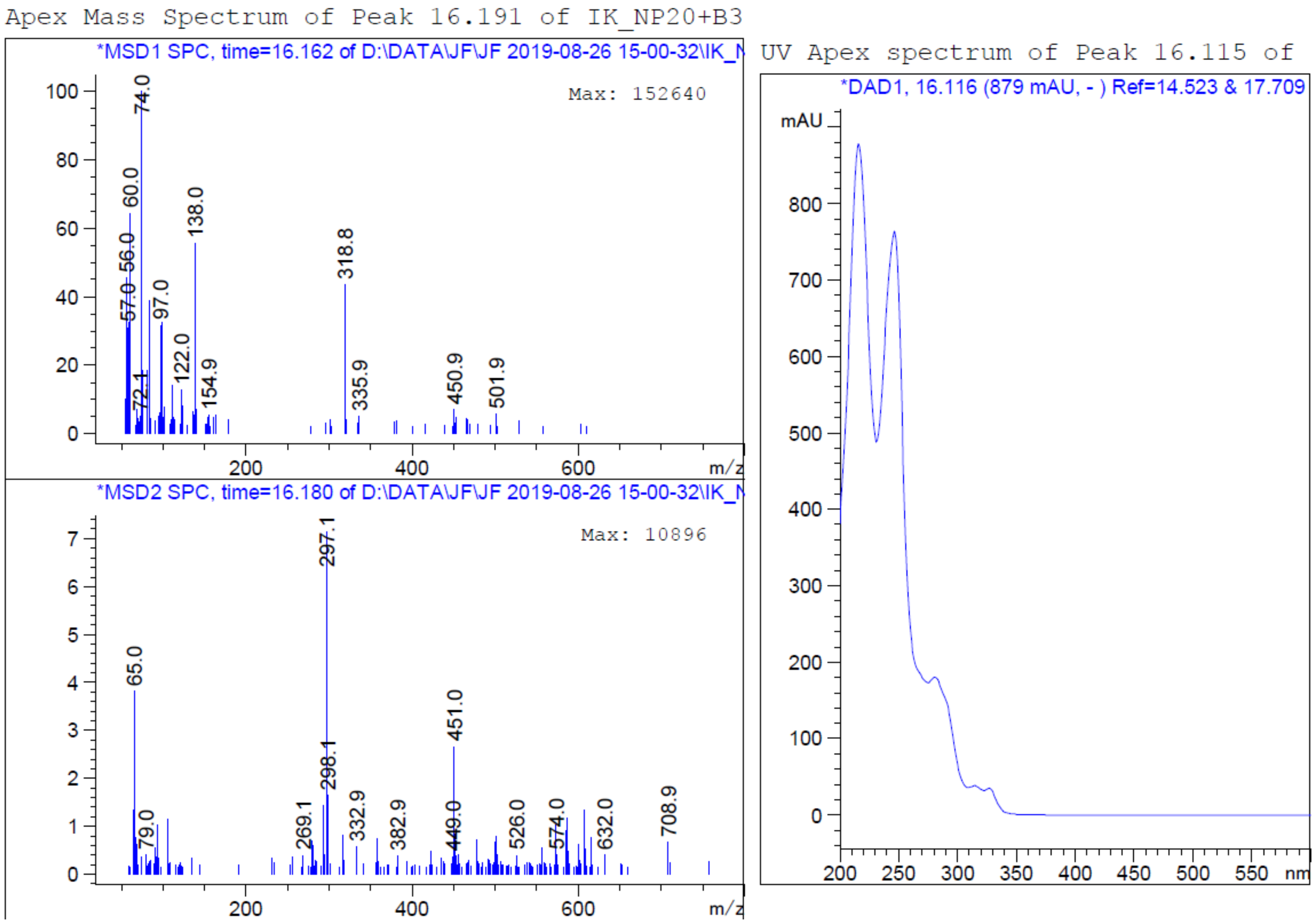


Mass spectra for fraction B3

**Fig. S4**. Mass spectra of the derived fractions from the toxic hydrophobic phase after fermentation of *Neofusicoccum parvum* Bt-67 with the plant surrender signal, ferulic acid. The figures represent the mass spectra information of the tested fractions; A5; A7; A8; A9, 10; B1, B2; B3 respectively.

### Fig. S5

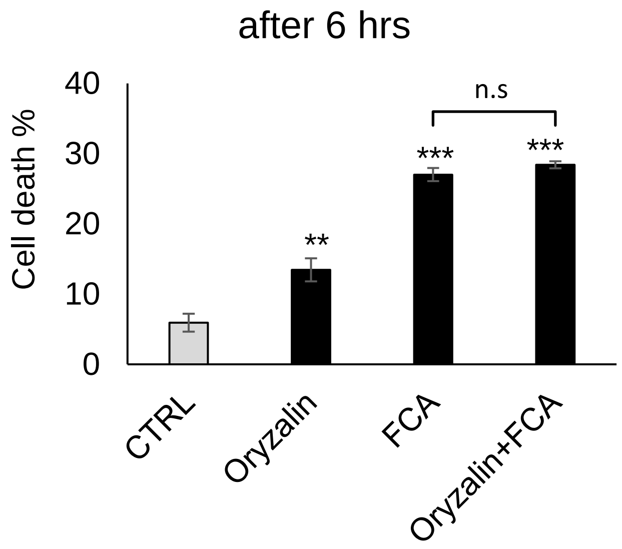


**Fig. S5.** Cell death rates by by 6 hrs of 6 µmol Fusicocin A after depletting the microtubules by 10 µmol Oryzalin. Asterisks and different letters indicate statistical differences based on LSD and Duncan's test with significant levels $P<0.05$ (*), $P<0.01$ (**), and $P<0.001$ (***)
