## Supplemental Method for "Hunting the plant surrender signal activating apoplexy in grapevines after *Neofusicoccum parvum* infection"

**The following supporting method is available for this article:**

**Methods S1.** **The effect of drought on fungal colonisation.**

To test the effect of drought stress on the infection development, infected grapevines of the variety *V. vinifera* cv. ‘Augster Weiss’ were grown under two water regimes. Each set consisted of infected plants and the respective mock control.The fungal DNA abundance was measured 30 days later in the infection site, but also at the lower and upper margin of the internode (3 cm below or above the infection site).

To evaluate the spread of the fungus through the internodes we quantified the fungal DNA after extracting genomic DNA from the wood as described in Cota-Sánchez, et al. (2006). We measured the abundance of fungal DNA by qPCR using 25 ng of DNA template, 1 unit of *Taq* polymerase (England Biolabs; Frankfurt, Germany), and specific *ITS-BOT* primers, which bind exclusively to *Botryosphaeriaceae* DNA, the primers sequence was as follows: (5’-AAACTCCAGTCAGTRAAC‘-3) as forward primer and (5’-TCCGAGGTCAMCCTTGAG‘-3) as reverse primers (Ridgway et al., 2011). The abundance of the fungal DNA was evaluated using a calibration curve (Flubacher, 2021), which was calculated based on a dilution series of transformed plasmid DNA amplified by TA cloning in the pGEM®-T Easy vector (ThermoFisher) according to the protocol of the producer.

**Extraction of fungal metabolites, HPLC and HPLC-MS for fractionation-guided activity**

The fungus was cultured in 20 L Yeast Malt Glucose medium (YMG, yeast extract 4.0 g^.^L^-1^, malt extract 10 g^.^L^-1^, glucose 10 g^.^L^-1^, the pH 5.5) in a fermenter (Braun, Melsungen, 120 rpm, 26°C and 3l/min aeration). After 5 days of fermentation (and a glucose level of 5 mM), we added 0.5 mM *trans*-ferulic acid to the culture. Afterwards the fermentation continued until full depletion of the free glucose. After separation of the culture fluid (18.5 L) from the mycelium by filtration (using a vacuum pump and 595 filter paper, Schleier & Schüll), we extracted the supernatant with ethyl acetate (12 L). After evaporation of the organic solvent to dryness under reduced pressure, we obtained 7.2 g of a crude extract, which we dissolved in MeOH to a concentration of 20 mg^.^mL^-1^. We used 20 µl from those concentrates for analysis by HPLC (Series 1100, Hewlett–Packard, Waldbronn, Germany; equipped with a LiChrospher RP18 column; 5 µm, 125x4 mm, Merck, Darmstadt, Germany). A gradient of 0.1% v/v formic acid and MeCN (method: 1% to 100% MeCN in 20 min; flow rate: 1 ml.min^-1^) served for fractionation into 96-well plates (Greiner Bio-One, Frickenhausen) for a bioactivity guided isolation as described by Buckel *et al*. (2013).

**References**:

**Buckel I, Molitor D, Liermann JC, Sandjo LP, Berkelmann-Löhnertz B, Opatz T, Thines E** (2013) Phytotoxic dioxolanone-type secondary metabolites from Guignardia bidwellii. Phytochemistry, 89, 96–103.

**Cota-Sánchez, JH, Remarchuk K, Ubayasena K** (2006) Ready-to-use DNA extracted with a CTAB method adapted for herbarium specimens and mucilaginous plant tissue. Plant Molecular Biology Reporter, 24(2), 161–167.

**Flubacher NS (2021).** 4-Hydroxyphenylacetic acid - a fungal polyketide suppressing plant defence. Dissertation, Karlsruhe Institute of Technology, Germany.

**Ridgway HJ, Amponsah NT, Brown DS, Baskarathevan J, Jones EE, Jaspers MV (2011**) Detection of botryosphaeriaceous species in environmental samples using a multi‐species primer pair. Plant Pathology, 60: 1118-1127.
