## Supplemental Table for "Hunting the plant surrender signal activating apoplexy in grapevines after *Neofusicoccum parvum* infection"

**The following supporting table is available for this article:**

**S. Table 1**: Genetic details and the primers sequences of the targeted genes:

| Gene | Accession nr. | Primer type | Primer sequence (5’-3’) | Reference |
| --- | --- | --- | --- | --- |
| VvEF1-α | XM_002284888 | VvEF1-α-F | GAACTGGGTGCTTGATAGGC | (Spagnolo et al., 2017) |
|  |  | VvEF1-α-R | AACCAAAATATCCGGAGTAAAAGA |  |
| VvUBQ | XR_002030723 | VvUBQ-F | GAGGGTCGTCAGGATTTGGA | Tina Moser (2014) |
|  |  | VvUBQ-R | GCCCTGCACTTACCATCTTTAAG |  |
| VvPR1 | XM002273752 | VvPR1-F | TGCTAACCAGAGATTGGCGATTG | Wang (2019) |
|  |  | VvPR1-R | CGCATCGGTGCCTGTCAATGAA |  |
| VvPAL | XM_002268220 | VvPAL-F | TCCTCCCGGAAAACAGCTG | (Spagnolo et al., 2017) |
|  |  | VvPAL-R | TCCTCCAAATGCCTCAAATCA |  |
| VvSTS27 | X76892 | VvSTS27-F | CCCAATGTGCCCACTTTAAT | Duan et al. (2015) |
|  |  | VvSTS27-R | CTGGGTGAGCAATCCAAAAT |  |
| VvSTS6 | XM_002271335 | VvSTS6-F | GTTGTGCTGCATAGCGTTGC | Vannozzi et al., (2012) |
|  |  | VvSTS6-R | GATTTAATTGGAAATTGTCCCCTTC |  |
| VvSTS16 | XM_019226034 | VvSTS16-F | CTTTTGACCCAATTGGAATCAAC | Vannozzi et al., (2012) |
|  |  | VvSTS16-R | TGACATGTTCCCATATTCACTTAG |  |
| VvSTS47 | AF274281 | VvRS1. F | TGGAAGCAACTAGGCATGTG | Duan et al. (2015) |
|  |  | VvRS1. R | GTGGCTTTTTCCCCCTTTAG |  |
| VvCAOMT | NM_001281171 | VvCAOMT-F | GGACTGGAGCCACCCTTAAC | This paper |
|  |  | VvCAOMT-R | CAACATGCTCCACACCAGGA |  |
| VvCAD | XM_002269320 | VvCAD-F | GAGAACGGTGAAGGGAAGCA | This paper |
|  |  | VvCAD-R | TCATTGTTTGCAAGGCGCTC |  |
| VvJAZ1 | JF900329 | VvJAZ1-F | TGCAGTCTGTTGAGCCAATACATA | Ismail et al., (2012) |
|  |  | VvJAZ1-R | CACGTTTCCGGACTTCTTTACAC |  |
| VvJAZ4 | XP_002272363.2. | VvJAZ4-F | TTCAGGAAATCGGCAACAACAGA | Zhang et al*.,* (2012) |
|  |  | VvJAZ4-R | CCCTTGGCGGCTAATAGCATG |  |
| VvJAZ9 | XP_002277157.1. | VvJAZ9-F | TTTACCGGGCAGAGAGCGCC | Zhang et al., (2012) |
|  |  | VvJAZ9-R | GATTCGGGCGTGCCGTTTCC |  |
| VvICS | XM019226638 | VvICS-F | CTCCGCCATCTCCCACTTGAAATC | Wang (2019) |
|  |  | VvICS-R | TCTTGTTGAGCGTGGAGCCAATC |  |
| VvCu.SOD1 | VIT_202s0025g04830.1 | VvCu.SOD1-F | GGAGCTCCTGACAGAGTTTATG | (Hu, et al., 2019) |
|  |  | VvCu.SOD1-R | ACCGAGAACCCTGACTACTT |  |
| VvCu.SOD2 | VIT_206s0061g00750.1 | VvCu.SOD2-F | CGACTGTCTCTGTTCGGATTAC | (Hu, et al., 2019) |
|  |  | VvCu.SOD2-R | GGATTGAAATGTGCTCCTGTTG |  |
| VvCu.SOD3 | VIT_208s0007g07280.1 | VvCu.SOD3-F | AGTGGGCAGCATTCCATT | (Hu, et al., 2019) |
|  |  | VvCu.SOD3-R | ACCAGCATTCCCAGTTGTT |  |
| Vv.Mn.SOD1 | VIT_00032675001 | VvMn.SOD1-F | CGGAGGTCATGTCAACCACT | This paper |
|  |  | VvMn.SOD1-R | ACCCAGTGAACCTTTTGGGG |  |
| Vv.Mn.SOD1 | VIT_213s0067g02990.1 | VvMn.SOD2-F | GGTGGTTGAAACTACTGCAAATC | (Hu, et al., 2019) |
|  |  | VvMn.SOD2-R | GTAATCCGGCCTCACATTCTT |  |

**References:**

**Duan D, Halter D, Baltenweck R, Tisch C, Tröster V, Kortekamp A, Hugueney P, Nick P** (2015) Genetic diversity of stilbene metabolism in Vitis sylvestris. Journal of Experimental Botany, 66(11), 3243–3257.

**Ismail A, Riemann M, Nick P** (2012) The jasmonate pathway mediates salt tolerance in grapevines. Journal of Experimental Botany, 63(5), 2127–2139.

**Hu X, Hao C, Cheng ZM, Zhong Y** (2019) Genome-wide identification, characterization, and expression analysis of the grapevine superoxide dismutase (SOD) family. International Journal of Genomics, 2019.

**Spagnolo A, Mondello V, Larignon P, Villaume S, Rabenoelina F, Clément C, Fontaine F** (2017) Defense responses in grapevine (Cv. Mourvèdre) after inoculation with the Botryosphaeria dieback pathogens Neofusicoccum parvum and Diplodia seriata and their relationship with flowering. International Journal of Molecular Sciences, 18(2), 1–12.

**Moser T** (2015) Investigation of the transcriptional regulation of candidate genes of pathogen defense against Plasmopara viticola in grapevines. Dissertationen, Julius Kühn-Institut, Germany.

**Vannozzi A, Dry IB, Fasoli M, Zenoni S, Lucchin M** (2012) Genome-wide analysis of the grapevine stilbene synthase multigenic family: genomic organization and expression profiles upon biotic and abiotic stresses. BMC Plant Biology, 12.

**Wang R** (2019) A New Actin-Dependent Pathway in Plant Defence -Aluminium Tolerance in Grapevine cells. PhD thesis. Karlsruhe Institute of Technology, Germany.

**Zhang Y, Gao M, Singer SD, Fei Z, Wang H, Wang X** (2012) Genome-Wide Identification and Analysis of the TIFY Gene Family in Grape. PLoS ONE, 7(9), 1–13.
